# DDX3 regulates the cap-independent translation of the Japanese encephalitis virus via its interactions with PABP1 and the untranslated regions of the viral genome

**DOI:** 10.1101/2024.10.15.618401

**Authors:** Chenxi Li, Linjie Zhang, Chenyang Tang, Xuan Chen, Jing Shi, Qingyu Li, Xue Jiao, Jinyao Guo, Bin Wang, Kefan Bu, Abudl Wahaab, Yuguo Yuan, Ming-an Sun, Yanhua Li

**Author notes:** Correspondence (Y.L.). These authors contributed equally to this work (Chenxi Li, Linjie Zhang).

## Abstract

The translation of global cellular proteins is almost completely repressed in cells with flavivirus infection, while viral translation remains efficient. The mechanisms of flaviviruses evade host translational shutoff are largely unknown. Here, we identified viral elements and host factors associated with JEV evasion of host shutoff. JEV 5′UTR lacked IRES or IRES-like activity, while noncapped 5′UTR initiated translation in the presence of 3′UTR. Furthermore, the elements DB2 and sHP-SL within 3′UTR were involved in the regulation of cap-independent translation, which is conserved in the genus *Orthoflavivirus*. By RNA affinity purification and mass spectrometry analysis, cellular DDX3 and PABP1 were identified as key factors in regulating cap-independent translation of JEV via their interactions with DB2 and sHP-SL RNA structures. Mechanistically, we revealed that DDX3 could bind to both 5′UTR and 3′UTR of the JEV genome to establish a closed-loop architecture, recruit eIF4G/eIF4A to form the DDX3/PABP1/eIF4G/eIF4A tetrameric complex via its interaction with PABP1, thereby recruiting 43S PIC to the 5′-end of the JEV genome to start translation. Our findings demonstrated a noncanonical translation strategy employed by JEV and further revealed the regulatory roles of DDX3 and PABP1 in this mechanism. These results expand our knowledge of the translation initiation regulation in flaviviruses under the state of host translational shutoff, which provides a conserved antiviral target against *orthoflavivirus*.

## Introduction

Japanese encephalitis virus (JEV) is a member of the genus *Orthoflavivirus* in the *Flaviviridae* family which includes many important zoonotic pathogens, such as Zika virus (ZIKV), Dengue virus (DENV), West Nile virus (WNV), and Tick-borne encephalitis virus (TBEV)(Erlanger et al., 2009; van Leur et al., 2021). The JEV genome comprises a single open reading frame (ORF) flanked by 5′ and 3′ untranslated regions (UTR). The 5′ UTR with a type I cap structure is around 100 nucleotides, while the 3′ UTR lacking a poly(A) tail is approximately 570 nucleotides. High-order RNA structures formed in 5′UTR and 3′UTR function as *cis*-acting elements required for viral RNA synthesis and viral protein translation(Zhang et al., 2022). JEV genome could serve as mRNA to translate a single polyprotein which is post-translationally processed by viral and host proteinases to generate three structural proteins (envelope [E], pre-membrane [prM], and capsid [C]) and seven nonstructural proteins (NS1, NS2A, NS2B, NS3, NS4A, NS4B, and NS5)(Chambers et al., 1990; Qiu et al., 2018). Based on the coding sequence of the E protein, JEV is phylogenetically classified into five genotypes (genotype I to V), and the prevalent strains mainly belong to genotype I (GI) and genotype Ⅲ (GⅢ) (Han et al., 2014; Li et al., 2020).

The translation initiation is the most important phase for the regulation of gene expression. In eukaryotes, the initiation of protein synthesis for most mRNAs generally occurs via the canonical cap-dependent mechanism(Gandin et al., 2022). To initiate cap-dependent translation, the eukaryotic initiation factor 4E (eIF4E) first recognizes and binds to the m^7^G(5′)ppp(5′)N cap structure at the 5′ end of mRNA, and then recruits an adaptor protein (eIF4G) and a RNA helicase complex (eIF4A and co-factor eIF4B) to form eIF4F cap-binding complex (eIF4E, eIF4G, and eIF4A), which further recruits the ribosomal 43S preinitiation complex (PIC) composed of a small 40S ribosomal subunit, the translation initiation factors (eIF1, eIF1A, eIF3, eIF5) and the ternary eIF2-GTP-Met-tRNAi complex, onto the mRNA(Sonenberg and Hinnebusch, 2009). Subsequently, the 43S PIC scans mRNAs in the 5′ to 3′ direction for the AUG start codon and recruits a 60S large ribosomal subunit to form an 80S monosome to initiate polypeptide synthesis(Jackson et al., 2010). Besides the necessary eIFs and ribosome subunits, various host proteins also participate in translation initiation. As a member of the DEAD (Asp-Glu-Ala-Asp) box family of RNA helicases, the multifunctional DEAD-box protein 3 (DDX3) contributes to the translation initiation process. DDX3 may accomplish modulation of cellular mRNA translation by its interactions with mRNA, poly(A)-binding protein 1 (PABP1), and initiation factors such as eIF2, eIF4E, and eIF4G(Geissler et al., 2012). In addition, DDX3 also interacts with the components of the 43S PIC, such as eIF3 and the 40S ribosomal subunit, thereby mediating the recruitment of 43S PIC to mRNAs(Lee et al., 2008; Soto-Rifo et al., 2012). Even though DDX3 was reported as a general translation initiation factor, DDX3 appears to only promote the translation of a subset of selected mRNAs with specific RNA structures at their 5′-end. Recently, DDX3 was demonstrated to enhance viral translation of JEV(Li et al., 2014), Hepatitis C virus (HCV) (Geissler et al., 2012), and Foot-and-mouth disease virus (FMDV)(Han et al., 2020) for efficient propagation. However, the mechanistic basis has not been fully understood.

Since viruses can not replicate outside of living cells, the expression of viral proteins exclusively relies on host translation apparatus. However, as an antiviral strategy, the cap-dependent translation in virus-infected eukaryotic cells is generally shut down to prevent the synthesis of viral proteins(Walsh et al., 2013; Walsh and Mohr, 2011). To overcome host shutoff or to compete for translational resources, viruses have evolved diverse mechanisms of cap-independent translation initiation to support robust expression of viral proteins. For instance, the VPg proteins of murine norovirus and feline calicivirus which serve as a substitute for the 5′ cap structure allow viral mRNA binding to eIF4E or even directly to eIF4G(Herbert et al., 1997; Royall et al., 2015). The internal ribosomal entry sites (IRESs) harbored by HCV(Yokoyama et al., 2019), bovine viral diarrhea virus (BVDV)(Burks et al., 2011), classical swine fever virus (CSFV)(Locker et al., 2007) and human immunodeficiency virus (HIV)(Plank et al., 2013), could directly recruit ribosomes to the internal region of viral mRNAs, omitting the need for eIF4E and eIF4G. Interestingly, several positive-strand RNA plant viruses, such as barley yellow dwarf virus(Bhardwaj et al., 2019), panicum mosaic virus(Chattopadhyay et al., 2014), and tomato bushy stunt virus(Fabian and White, 2004), could employ a cap-independent translation element (CITE) residing in the 3′UTR to initiate translation. This mechanism of viral translation initiates at the 5′-end with the assistance of viral 3′UTR. Notably, the cap-dependent and cap-independent translations are not mutually exclusive and may synergically enhance viral translation.

In response to cellular stress triggered by flavivirus infection, cellular protein translation is often controlled at the rate-limiting step of initiation(Roth et al., 2017; Tu et al., 2012; Wang et al., 2020). The repression of the initiation step of cap-dependent translation was documented for DENV, WNV, TBEV, TMUV, and ZIKV(Roth et al., 2017; Wang et al., 2020). Despite host translational shutoff, the synthesis of flavivirus proteins remains efficient in supporting viral replication. DENV employs a novel non-IRES-mediated non-canonical translation mechanism that requires the interaction between DENV 5′UTR and 3′UTR(Edgil et al., 2006), while the 5′UTR of ZIKV without a type-I cap structure harbored IRES activity which directly initiates cap-independent translation(Song et al., 2019). In addition, *cis*-acting elements in TMUV 3′UTR were shown to play crucial regulatory roles in cap-independent translation(Wang et al., 2020). Thus, flaviviruses might have various strategies to regulate non-canonical translation initiation. Of note, experimental evidence supporting the non-canonical translation of JEV is lacking. In this study, we found that JEV can be rescued using genomic RNA without a cap structure via cap-independent translation in cells of different species. JEV 5′UTR lacks IRES or IRES-like activity, while 5′UTR could initiate the cap-independent translation in the presence of 3′UTR. Mechanistically, we revealed that DB2 and sHP-SL elements within 3′UTR play decisive roles in the regulation of cap-independent translation via their interactions with DDX3 and PABP1. As a translation initiation factor, DDX3 anchors to both 5 ′ UTR and 3′UTR of the JEV genome, forming the DDX3/PABP1/eIF4G/eIF4A tetrameric complex via its interaction with PABP1, thereby recruiting the 43S PIC to the 5′-end of the JEV genome and allowing translation initiation. These results provide new insights for the understanding of the translational initiation regulation of flaviviruses.

## Results

### JEV adopts a cap-independent translation strategy to evade host shutoff

Since their genomes harbor a type I cap structure at the 5′end, flaviviruses are supposed to initiate viral translation via the cap-dependent mechanism(Barrows et al., 2018). However, the synthesis of viral polyproteins remains efficient when the cap-dependent host cellular translation is suppressed at the level of translation initiation during flavivirus infection(Barrows et al., 2018; Roth et al., 2017). Here, host translational shutoff induced by JEV infection was confirmed in various cell lines using a puromycin incorporation assay (Figure 1-figure supplement 1A-1D). In contrast, the expression level of NS1′ gradually increased as infection went on, suggesting that JEV translation could evade the shutoff of cap-dependent translation initiation. In BHK-21 and ST cells, downregulation of cap-dependent translation by reduction of eIF4E expression or interfering with the interaction between eIF4E and eIF4G using the inhibitor 4E2RCat had no obvious effect on NS1′ expression and viral yields (Figure 1A-1D and Figure 1-figure supplement 1E-1H), suggesting that JEV translation could be initiated with other mechanisms. To further confirm whether type I cap structure is indispensable for the translation initiation and virus recovery of JEV, three types of genomic RNA of the GI JEV with different 5′ ends illustrated in Figure 1E were synthesized for virus recovery. The JEV RNA transcripts were respectively transfected into four cell lines derived from different host species. An obvious cytopathic effect (CPE) characterized by cell shrinkage, rounding, necrosis, and detachment was observed in all cell lines except C6/36 cells (Figure 1F). Meanwhile, the NS1′ expression detected by IFA confirmed that recombinant viruses were successfully rescued in all cell lines using three types of JEV genomic RNAs, even though CPE was not observed in C6/36 cells (Figure 1F). Further, the yields of infectious virus in culture supernatants were monitored at 24, 36 and 48 hours post-transfection (hpt). Three JEV genomic RNAs produced similar viral titers in BHK-21 and C6/36 cells at all time points(Figure 1G), which was consistent with previous studies on other flaviviruses(Song et al., 2019). However, viral titers produced with the noncapped genomic RNA were significantly lower than those of the capped genomic RNAs in ST and DF-1 cells(Figure 1G). These results suggested that a cap-independent translation initiation strategy could be employed by JEV to evade host shutoff.

**Figure 1.**
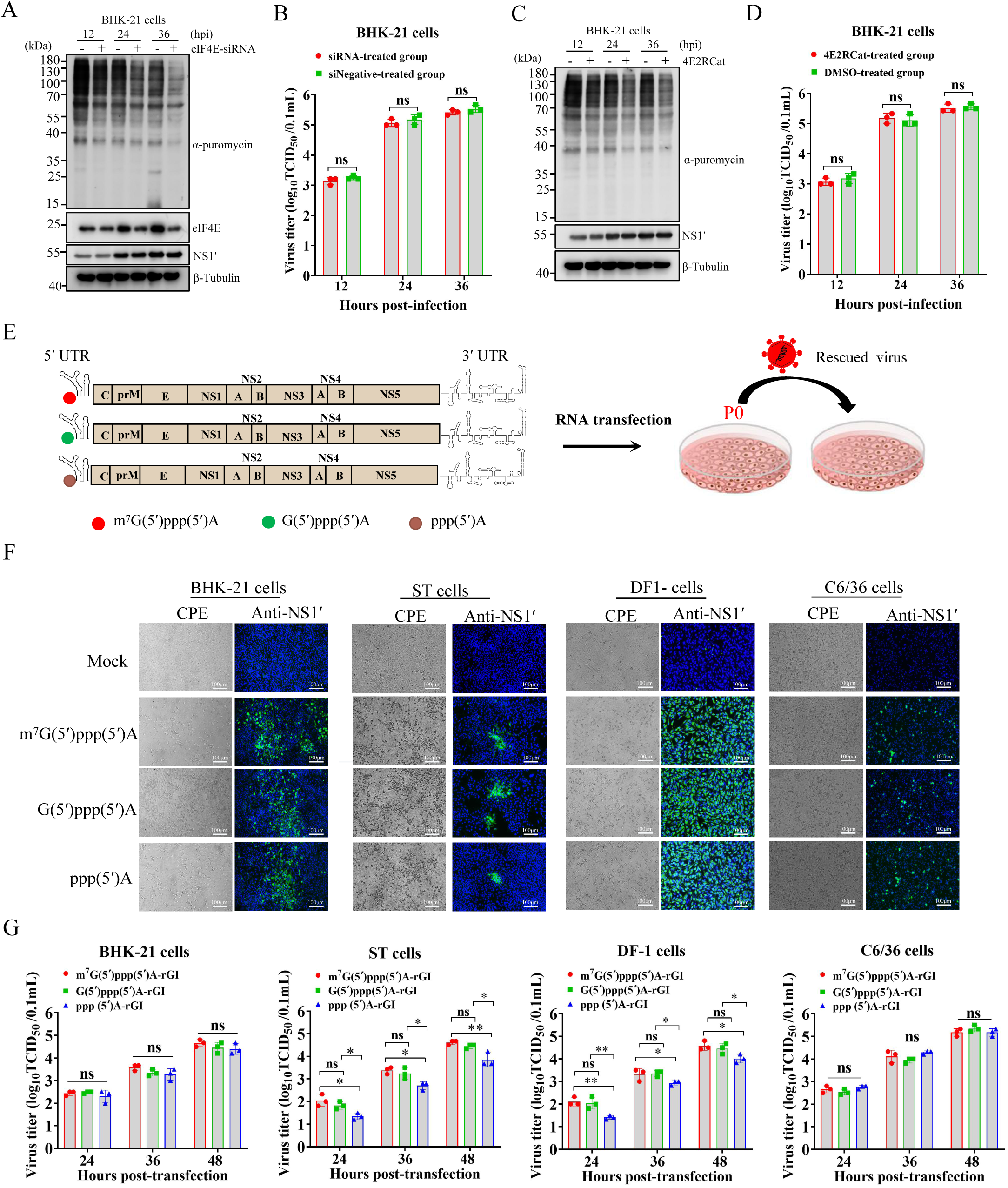
Both cap-dependent and cap-independent translation initiation strategies are involved in the expression of JEV proteins. (A-D) BHK-21 cells were respectively transfected with 100 pmol of a mixture of eIF4E-specific siRNAs (Table supplement 3) or treated with 20 μM 4E2RCat for 12 hours, and then infected with JEV at an MOI of 0.1. At different time points of post-infection, the cells were labeled with puromycin for 30 min and harvested to analyze puromycin incorporation (A and C). The viral titers in culture supernatants of BHK-21 cells treated with eIF4E-specific siRNAs (B) or 4E2RCat (D) were measured by TCID_50_ assay (n=3). (E) Schematic diagram of the virus rescue using JEV RNA transcripts modified with three different 5′ termini: m7G(5′)ppp(5′)A, G(5′)ppp(5′)A and ppp(5′)A. (F) Cytopathic effects and immunofluorescence assay of cells transfected with full-length viral RNA transcripts with different 5′ termini. (G) The viral titers in culture supernatants of cells transfected with 2μg RNA transcripts at 24, 36 and 48hpt (n=3). *, p<0.05; **, p<0.01; ns, no statistical difference. Data are presented as mean ± standard deviation (SD) of three independent experiments and tested by Student’s *t*-test (B, D, and G).

### RNA structures in UTRs play essential roles in cap-independent translation initiation

In most single-stranded positive-sense RNA viruses without a cap structure at the 5′ end of their genomes, the IRES element situated at the 5′ end of the viral genomes is usually employed to initiate viral translation. Several secondary RNA structures at the 5′ end of the JEV genome are required for viral translation and RNA synthesis, including SLA, SLB, and cHP (Figure 2A)(Chiu et al., 2005; Upstone et al., 2023). A panel of bicistronic reporter plasmids as depicted in Figure 2-figure supplement 1A was generated to evaluate the IRES or IRES-like activity of the 5′-end of the JEV genome. In those constructs, *Renilla* luciferase (RLuc) is expressed through cap-dependent translation, while *firefly* luciferase (Fluc) is expressed dependent on the IRES or IRES-like activity of the upstream viral sequences. HCV IRES (HCV-5′ UTR) and inactivated HCV IRES (HCV-5′ UTR-Δdomain III) were used as positive and negative controls. In DNA transfected BHK-21 cells all constructs expressed similar high levels of RLuc, while neither 5′UTR nor 5′UTR-cHP-cCS effectively initiated the cap-independent translation of FLuc in comparison to a significantly higher level of FLuc expression driven by HCV IRES (Figure 2B). To rule out the possibility that the nonviral sequence added to the 5′ terminus may interfere the IRES activity of JEV 5′UTR, a panel of monocistronic RNA reporters without a cap structure, including JEV-5′UTR-FLuc, JEV 5′UTR-cHP-cCS-FLuc, HCV-5′UTR-FLuc, and HCV-5′UTR-Δdomain III-FLuc (Figure 2-figure supplement 1B), were transfected into BHK-21 cells. In line with results generated with the bicistronic luciferase assay, the JEV-5′UTR-FLuc and JEV 5′UTR-cHP-cCS-FLuc mRNA produced similar low levels of FLuc as the inactivated HCV IRES, while HCV-5′UTR-FLuc mRNA produced a significantly higher level of FLuc (Figure 2C). Thus, the 5′UTR or 5′UTR-cHP-cCS of JEV does not have IRES or IRES-like activity.

**Figure 2.**
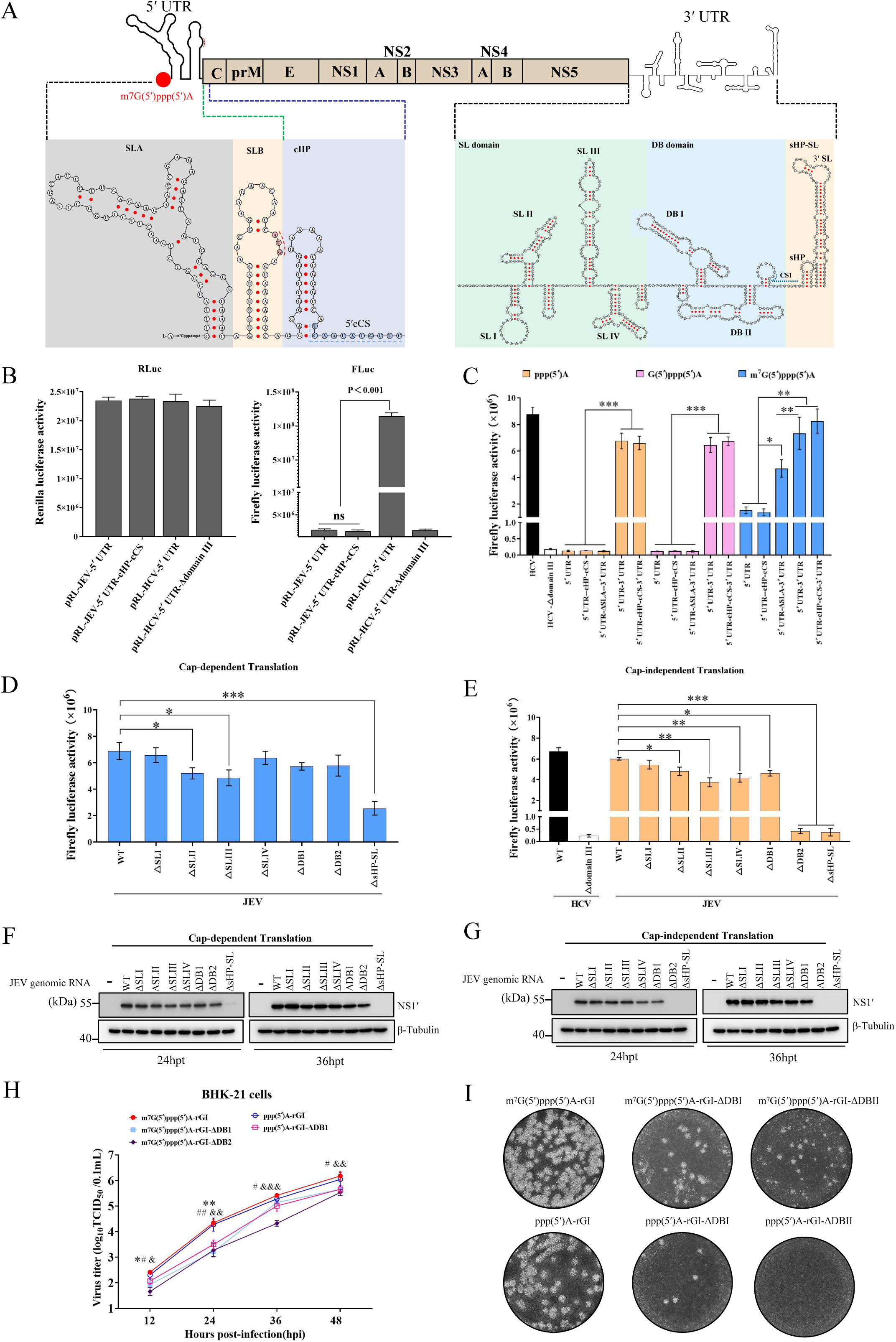
Three cis-acting elements in UTRs are crucial for the cap-independent translation initiation of JEV mRNA. (A) Secondary structure diagram of the 5′- and 3′- termini of JEV. (B) BHK-21 cells were respectively transfected with bicistronic constructs pRL-JEV-5′UTR, pRL-JEV-5′UTR-cHP-cCS, pRL-HCV-5′UTR and pRL-HCV-5′UTR-Δdomain III at a dose of 2 μg. At 24 hours post-transfection, the *firefly* luciferase and *Renilla* luciferase activities in BHK-21 cells were determined by luciferase assay (n=4). (C-E) Monocistronic reporter construct or its deletion mutants were generated via T7 promoter-mediated *in vitro* transcription, and then transfected into BHK-21 cells. At 12 hours post-transfection, the *firefly* luciferase activity in BHK-21 cells was determined by luciferase assay (n=4; *, p<0.05, **, p<0.01, ***, p<0.001, ns, no significance; statistical significance determined by one-way ANOVA). (F and G) Western-blot analysis of BHK-21 cells transfected with the JEV genomic RNA with 5′termini m^7^G(5′)ppp(5′)A or ppp(5′)A of WT or deletion mutants. (H) BHK cells were infected with viruses m^7^Gppp(5′)A-rGI, ppp(5′)A-rGI, m^7^G(5′)ppp(5′)A-rGI-ΔDB1, ppp(5′)A-rGI-ΔDB1 and m^7^G(5′)ppp(5′)A-rGI-ΔDB2 at an MOI of 0.05. At the indicated time points, the culture supernatants were collected to determine virus titers by TCID_50_ assay (n=3). Data are the means ± SD of three or four independent experiments, and statistical significance was tested by one-way ANOVA analysis with Tukey’s multiple comparison test. The significant differences between m^7^Gppp(5′)A-rGI and m^7^G(5′)ppp(5′)A-rGI-ΔDB1 are labeled (*, p<0.05; **, p<0.01). The significant differences between m^7^Gppp(5′)A-rGI and ppp(5′)A-rGI-ΔDB1 is marked (**^#^**, p<0.05; **^##^**,p<0.01). The significant differences between m^7^Gppp(5′)A-rGI and m^7^G(5′)ppp(5′)A-rGI-ΔDB2 is indicated (^&^, p<0.05; ^&&^, p<0.01; ^&&&^, p<0.001). (I) Plaque morphology of the recombinant viruses in BHK-21 cells.

The non-polyadenylated 3′UTR of flavivirus serves as an enhancer of viral translation initiation and translation efficiency(Berzal-Herranz et al., 2022; Holden and Harris, 2004). To assess the role of JEV 3′UTR in cap-independent translation, a panel of RNA transcripts with JEV 3′UTR following the FLuc ORF was designed (Figure 2-figure supplement 1B). Three types of RNA transcripts with or without a functional cap structure were synthesized for each reporter. In RNA transfected BHK-21 cells, all transcripts with a functional cap structure produced high levels of FLuc, and the capped RNA transcripts with 3′UTR produced nearly 5-fold more FLuc than the capped RNA transcripts without 3′UTR (Figure 2C), confirming that flavivirus 3′UTR contributes to the enhancement of viral translation(Berzal-Herranz et al., 2022). Of note, in the presence of 3′UTR, the RNA transcripts without a functional cap structure produced comparable high levels of FLuc as the capped RNA transcripts, implying the involvement of JEV 3′UTR in the regulation of cap-independent translation initiation. Moreover, the absence of cHP-cCS in JEV-5′UTR-FLuc-3′UTR did not affect the expression of FLuc, suggesting that cHP and cCS are not important for cap-independent translation (Figure 2C). Noteworthy, the essential role of the SLA structure of 5′UTR in cap-independent translation was proved by the extremely low FLuc produced by the reporter JEV-5′UTR-ΔSLA-FLuc-3′UTR with a functional cap (Figure 2C), indicating that the integrity of JEV 5′UTR structure is also crucial for cap-independent translation. JEV 3′UTR forms high-order RNA structures, including SLⅠ, SLⅡ, SLⅢ, SLⅣ, DB1, DB2, and sHP-SL (Figure 2A)(Alvarez et al., 2005; Zhang et al., 2022). To evaluate the role of each motif of 3′UTR in cap-independent translation, a panel of JEV-5′UTR-FLuc-3′UTR mutants was generated with the individual motif deleted (Figure 2-figure supplement 1C). All capped RNA transcripts effectively initiated translation in a cap-dependent manner, although some mutants produced relatively low FLuc compared to the JEV-5′UTR-FLuc-3′UTR (Figure 2D). For the cap-independent translation using the noncapped RNA transcripts, the deletion of each motif downregulated FLuc expression and the deletion of DB2 or sHP-SL reduced FLuc expression to background level as the negative control HCV-5′UTR-Δdomain III-FLuc (Figure 2E), suggesting that DB2 and sHP-SL are critical for JEV cap-independent translation initiation.

Next, a panel of mutants with the individual motif of 3′UTR deleted was created with a reverse genetic system of GI JEV Beijing/2020-1(Li et al., 2023) to confirm the crucial RNA structures of 3′UTR in the context of JEV infection. The full-length genomic RNAs with or without a functional type I cap were respectively transfected BHK-21 cells to rescue recombinant viruses. All RNA mutants, except for ΔsHP-SL, successfully expressed viral protein and rescued recombinant viruses in a cap-dependent manner (Figure 2F and Figure 2-figure supplement 2A), while the noncapped RNAs of ΔDB2 and ΔsHP-SL failed to express viral protein and rescue viruses (Figure 2G and Figure 2-figure supplement 2A). These results confirmed the indispensable role of DB2 in initiating cap-independent translation. The designed deletions of the rescued viruses (P0) were validated by Sanger sequencing (Figure 2-figure supplement 3). All mutants with individual SL structure deleted exhibited comparable growth kinetics as the WT virus in BHK-21 cells (Figure 2-figure supplement 2B and 2C), while the mutants of ΔDB1 and ΔDB2 produced viral titers significantly lower than those of the WT virus (Figure 2H). Analysis of plaque sizes formed in BHK-21 cells indicated that mutant viruses of ΔDB1 and ΔDB2 formed small plaques approximately 2-fold smaller than those formed by the WT virus (Figure 2I).

Of note, both capped and noncapped JEV genomes without sHP-SL failed to rescue viable viruses. Considering that the cyclization sequence (3′CS) within sHP-SL is indispensable for viral RNA synthesis and virion production(Yu and Markoff, 2005; Yun et al., 2009), we explored whether the failure of ΔsHP-SL mutant recovery was caused by a defect of viral RNA synthesis. A JEV replicon (rGI-GLuc) as depicted in Figure 2-figure supplement 4 allows discrimination between viral translation and viral RNA synthesis, and a replication-defective RNA with NS5-372-386 deletion (GI-GLuc-NS5mut) was included as a negative control. Viral RNA synthesis could be assessed based on the second round of GLuc accumulation for rGI-GLuc but not for rGI-GLuc-NS5mut (Figure 2-figure supplement 4B). The deletions of DB2 and sHP-SL completely blocked GLuc expression in a cap-independent manner, while the deletion of sHP-SL also downregulated the cap-dependent translation of GLuc (Figure 2-figure supplement 4C and 4D). Based on GLuc expression at 8∼12 hpt, the replicon without sHP-SL is replication-defective, and the deletion of DB2 also attenuated viral RNA synthesis (Figure 2-figure supplement 4C and 4D). Thus, DB2 and sHP-SL within 3′UTR are essential for JEV cap-independent translation and are also involved in viral RNA synthesis.

### DB2 and sHP-SL determine JEV evasion of host translational shutoff and are associated with the virulence of JEV in mice

The role of cap-independent translation initiation in JEV evasion of host translational shutoff was initially studied with the monocistronic reporters in Figure 2-figure supplement 1B. To validate the critical role of DB2 and sHP-SL in JEV evasion of host translational shutoff, we first evaluated the translational activity of luciferase reporter RNA in the presence of suppression of cap-dependent translation. The suppression of cap-dependent translation initiation significantly reduced FLuc expression of reporter RNAs without DB2 or sHP-SL in a dose-dependent manner but not that of WT (Figure 3A and 3B). Furthermore, the suppression of cap-dependent translation resulted in a decrease in the translational efficiency of viral RNA without DB2 or sHP-SL (Figure 3C-3E, and Figure 3-figure supplement 1), and extremely low viral titers with the capped ΔDB2 RNA at all time points (Figure 3D-3F), suggesting that both DB2 and sHP-SL are critical for JEV evasion of the shutoff of cap-dependent translation. Meanwhile, the effect of suppressing cap-dependent translation initiation on the growth kinetics of rGI and rGI-ΔDB2 was further evaluated. Similar growth kinetics of rGI were observed in BHK-21 cells with/without cap-dependent translation attenuated (Figure 3G and 3I). In contrast, eIF4E knockdown and 4E2RCat treatment significantly attenuated the growth of rGI-ΔDB2 (Figure 3G and 3I). Consistently, the expression of NS1′ was downregulated by eIF4E knockdown and 4E2RCat treatment for rGI-ΔDB2 but not WT virus (Figure 3H and 3J). Thus, DB2 and sHP-SL are critical for JEV evasion of host translational shutoff.

**Figure 3.**
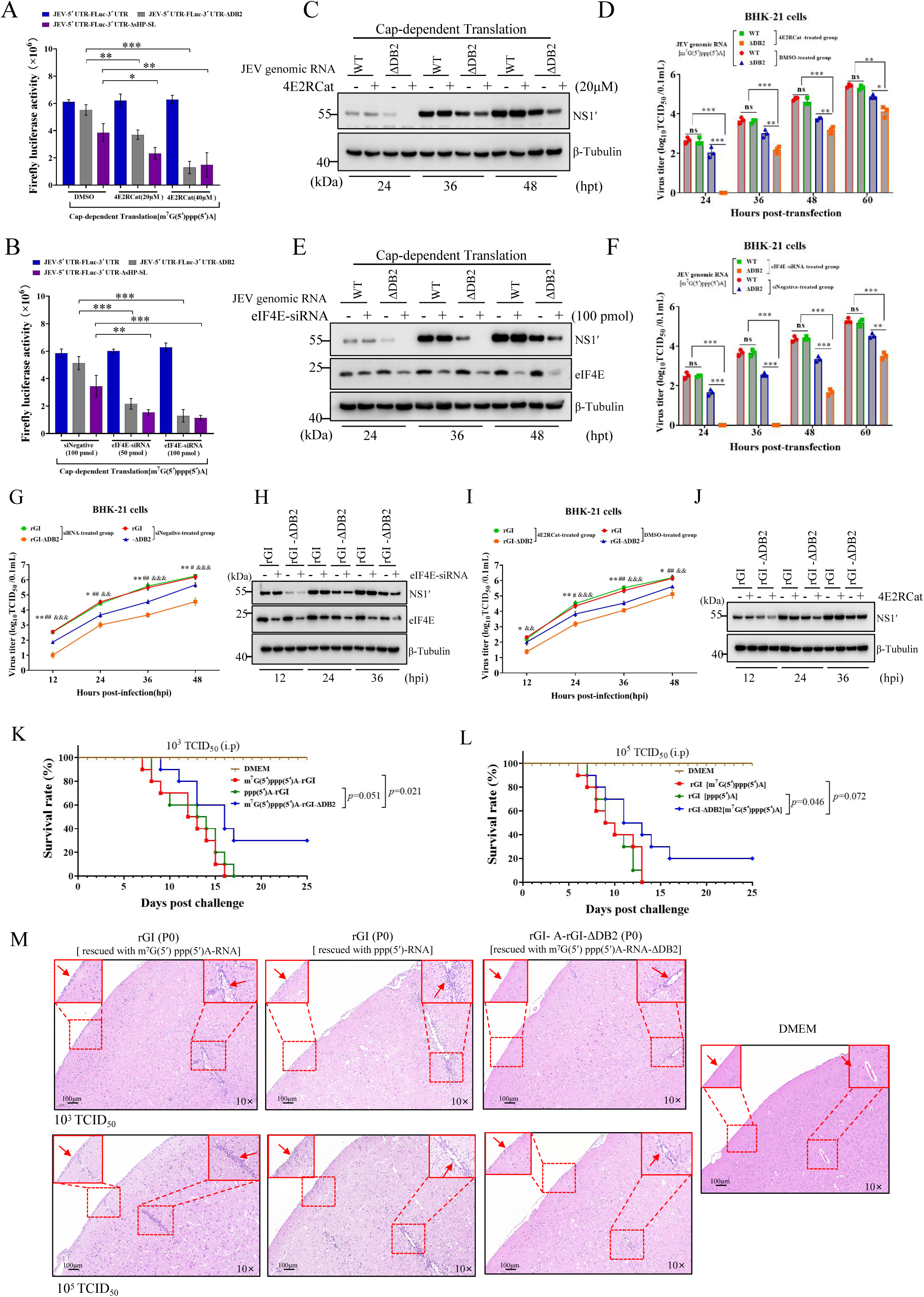
The *cis*-acting elements DB2 and sHP-SL of 3′UTR determine the resistance of JEV to the suppression of cap-dependent translation initiation and the virulence of JEV in mice. (A and B) The *firefly* luciferase activity assay of monocistronic RNA reporters with 5’terminal modified by either m^7^G(5′)ppp(5′)A or ppp(5′) in BHK-21 cells treated with eIF4E-specific siRNAs or 4E2RCat (n=4). (C-F) BHK-21 cells were respectively treated with eIF4E-specific siRNAs (100pmol) or 4E2RCat (20μM) and then transfected with JEV genomic RNA of WT or ΔDB2 mutant with a type I cap structure. At the indicated time points of post-transfection, NS1′ protein was detected by immunoblotting (C and E) and viral titers in culture supernatants were measured by TCID_50_ assays (D and F) (n=3). (G-J) BHK-21 cells were respectively treated with eIF4E-specific siRNAs or 4E2RCat and then infected with rGI and rGI-ΔDB2 at an MOI of 0.05. At the indicated time points of post-infection, viral titers in culture supernatants were measured by TCID_50_ assays (G and I) (n=3), and NS1′ protein was detected by immunoblotting (H and J). (K and L) The survival rate of mice (n=10) intraperitoneally mock-infected or infected with the indicated JEV at doses of 10^3^ and 10^5^ TCID_50_. The significant differences were determined by the Kaplan-Meier analysis. (M) Histopathological analysis of brain lesions of the dead mice. *, p<0.05; **, p<0.01; ***, p<0.001. Data are presented as mean ± SD and tested by one-way ANOVA (A-B, D, F, G and I).

Next, the effect of DB2 deletion on the pathogenicity of JEV was evaluated via mice challenge. All mice succumbed to challenges at different doses in the rGI groups, while 20%∼30% survival rates were observed in the rGI-ΔDB2 groups (Figure 3K and 3L). In the brains of dead mice, viral RNA was detected by RT-PCR, and the DB2 deletion was verified. The characteristic lesion of JEV encephalitis was observed in the brain of all dead mice, but it was reduced in the rGI-ΔDB2 group (Figure 3M). These results suggested that DB2 is associated with the virulence of JEV in mice. Due to the lethal effect of sHP-SL deletion on JEV, it is not possible to assess the role played by sHP-SL in JEV pathogenicity.

### The critical role of DB2 and sHP-SL in the cap-independent translation is highly conserved in the genus *orthoflavivirus*

Cap-independent translation initiation has been proven in several flaviviruses(Edgil et al., 2006; Song et al., 2019; Wang et al., 2020). Based on in silico prediction, the secondary structures of DB2 and sHP-SL are highly conserved among different JEV genotypes, ZIKV, WNV, DENV and TMUV (Figure 4A). We speculated that the function of DB2 and sHP-SL in regulating cap-independent translation is conserved in flaviviruses. A panel of monocistronic FLuc reporters carrying different flavivirus UTRs with/without the deletion of DB2 or sHP-SL was utilized to evaluate the function of flavivirus UTRs in the cap-independent translation (Figure 4B). As expected, all noncapped reporter RNAs flanked by 5′UTR of JEV and 3′UTR of ZIKV, TMUV, or various genotypes of JEV produced comparable levels of FLuc as the corresponding capped RNAs (Figure 4C). Noteworthy, all reporter RNAs with the absence of either DB2 or sHP-SL produced extremely low levels of FLuc, suggesting that DB2 and sHP-SL are critical for cap-independent translation initiation in various genotypes of JEV and other flaviviruses. We also confirmed these results in the context of virus recovery of GⅢ and GⅤ JEV. As illustrated in Figure 4D, infectious cDNA clones of GⅢ and chimeric GⅠ/GⅤ-UTR were used to create mutants with a deletion of DB2 or sHP-SL. The capped and noncapped genomic RNAs were transfected into BHK-21 cells to rescue recombinant viruses. All WT genomes successfully produced infectious viruses, while the absence of DB2 and sHP-SL prevented three JEV genotypes from the cap-independent translation leading to the failure of virus recovery (Figure 4E and F). These results further supported that DB2 and sHP-SL are essential to the cap-independent translation in various genotypes of JEV. Taken together, the decisive role of DB2 and sHP-SL in regulating cap-independent translation is evolutionarily conserved in various flaviviruses.

**Figure 4.**
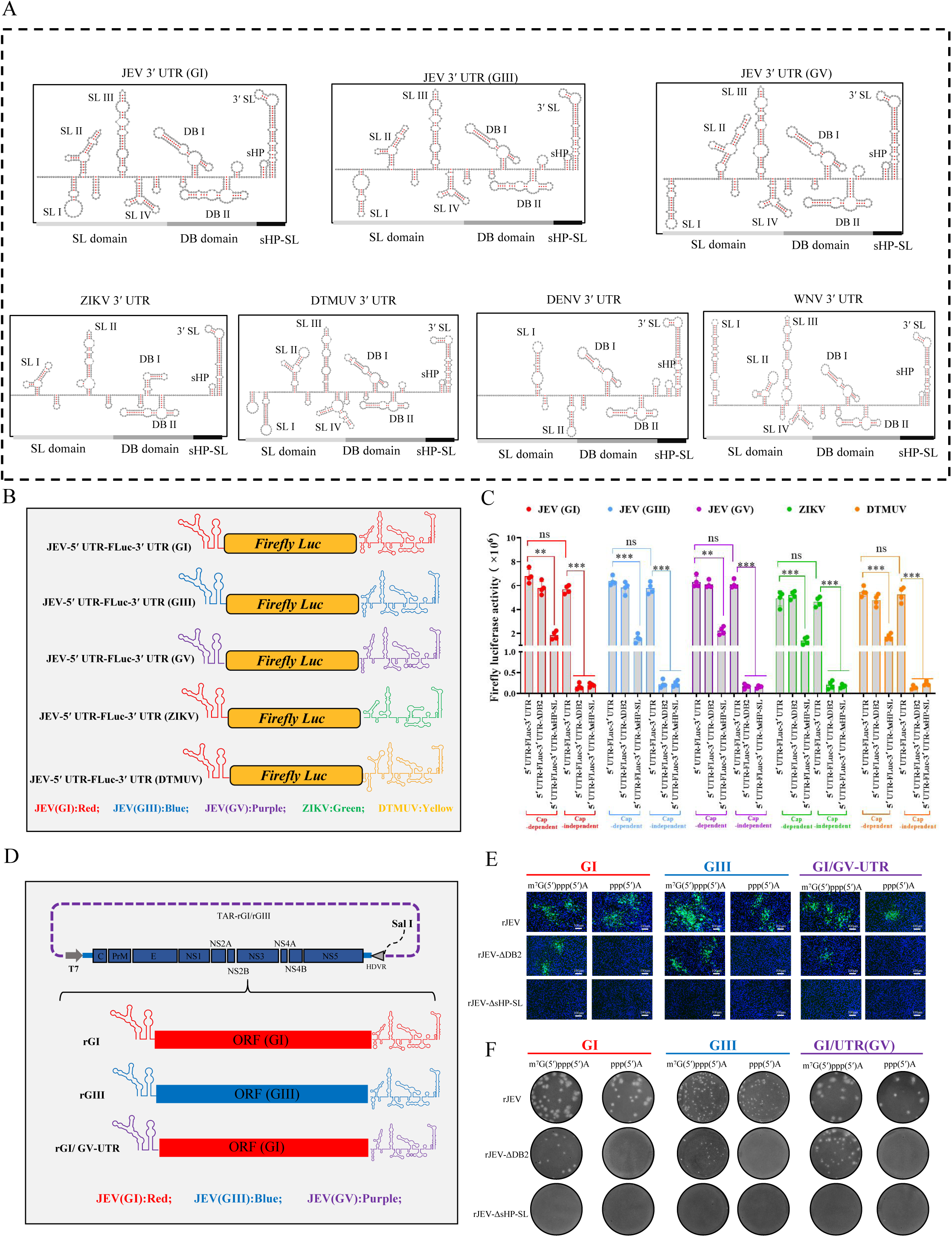
The critical role of DB2 and sHP-SL within 3′UTR in cap-independent translation initiation is evolutionarily conserved in various flaviviruses. (A) Secondary structure diagram of 3′UTR of various genotypes JEV, ZIKV, DENV, WNV and TMUV. (B) Diagrams of flaviviruses monocistronic reporter constructs controlled by T7 promoter: JEV-5′UTR-FLuc-3′UTR(GI), JEV-5′UTR-FLuc-3′UTR (GIII), JEV-5′UTR-FLuc-3′UTR (GV), ZIKV-5′ UTR-FLuc-3′ UTR and TMUV-5′UTR-FLuc-3′UTR. (C) Monocistronic RNA reporters with 5’terminal modified by either m^7^G(5′)ppp(5′)A or ppp(5′) were generated via T7 promoter-mediated *in vitro* transcription, and then transfected into BHK-21 cells. At 12 hours post-transfection, the cell samples were harvested for *firefly* luciferase activity assay (n=4). ***, p<0.001; **, p<0.01; ns, no significance, tested by the one-way ANOVA analysis with Tukey’s multiple comparison test(C). Data are the means ± SD of results of four independent experiments. (D) Schematic illustration of constructed infectious clones of GI JEV, GIII JEV and GI/GV-UTR JEV. (E and F) Immunofluorescence analysis and plaque morphologies of rescued viruses rJEV, rJEV-ΔDB2 and rJEV-ΔsHP-SL in BHK-21 cells.

### Cellular DDX3 and PABP1 are involved in JEV cap-independent translation

As obligate intracellular parasites, viruses must exploit the host translation apparatus to support viral protein synthesis. The interactions between viral UTR and cellular factors are believed to be the first step of viral translation initiation. A biotinylated RNA pull-down assay was performed to identify host proteins associated with DB2 and sHP-SL of JEV. As shown in Figure 5A, a unique band specifically associated with the biotinylated JEV 3′UTR instead of biotinylated JEV 3′UTR-ΔDB2-sHP-SL was observed for two cell lines. LC-MS/MS analysis with this band identified 16 proteins for BHK-21 cells and 21 proteins for 293T cells, with six proteins shared by the two cell lines (Figure 5A, Figure 5-figure supplement 1, and Table supplement 1). The shared six proteins were respectively knocked down using siRNA in BHK-21 and 293T cells to test whether they play roles in cap-independent translation. The cap-independent FLuc translation of JEV-5′UTR-FLuc-3′UTR was significantly reduced only when DDX3 or PABP1 was knocked down (Figure 5B). The siRNA knockdown of DDX3 did not affect the cap-dependent FLuc expression, while PABP1 knockdown reduced 60% of cap-dependent translation (Figure 5C). Consistently, the overexpression of DDX3 specifically enhanced cap-independent FLuc expression, while the ectopically expressed PABP1 promoted both cap-dependent and cap-independent FLuc expression (Figure 5D). To further strengthen the authentic roles of DDX3 and PABP1 in regulating cap-independent translation of JEV, the translational activity of viral genomic RNA was tested in DDX3 or PABP1 knock-down BHK-21 cells. The siRNA knockdown of DDX3 significantly reduced NS1′ protein expression and viral production of noncapped WT JEV genomes (Figure 5E and 5F). Besides downregulating the translational activity of noncapped WT RNA, DDX3 knockdown also obviously weakened NS1′ protein expression and viral production of capped WT RNA, but did not affect those of capped ΔDB2 mutant RNA that is defective in cap-independent translation (Figure 5E and 5F). Of note, PABP1 knockdown reduced both the cap-dependent translation of WT RNA and Δ DB2 mutant RNA and the cap-independent translation of WT RNA (Figure 5G and 5H), which is consistent with the results using FLuc reporters (Figure 5C), indicating that PABP1 is required for both cap-dependent and cap-independent translations of JEV mRNA. Thus, we confirmed that both cellular DDX3 and PABP1 are involved in JEV cap-independent translation.

**Figure 5.**
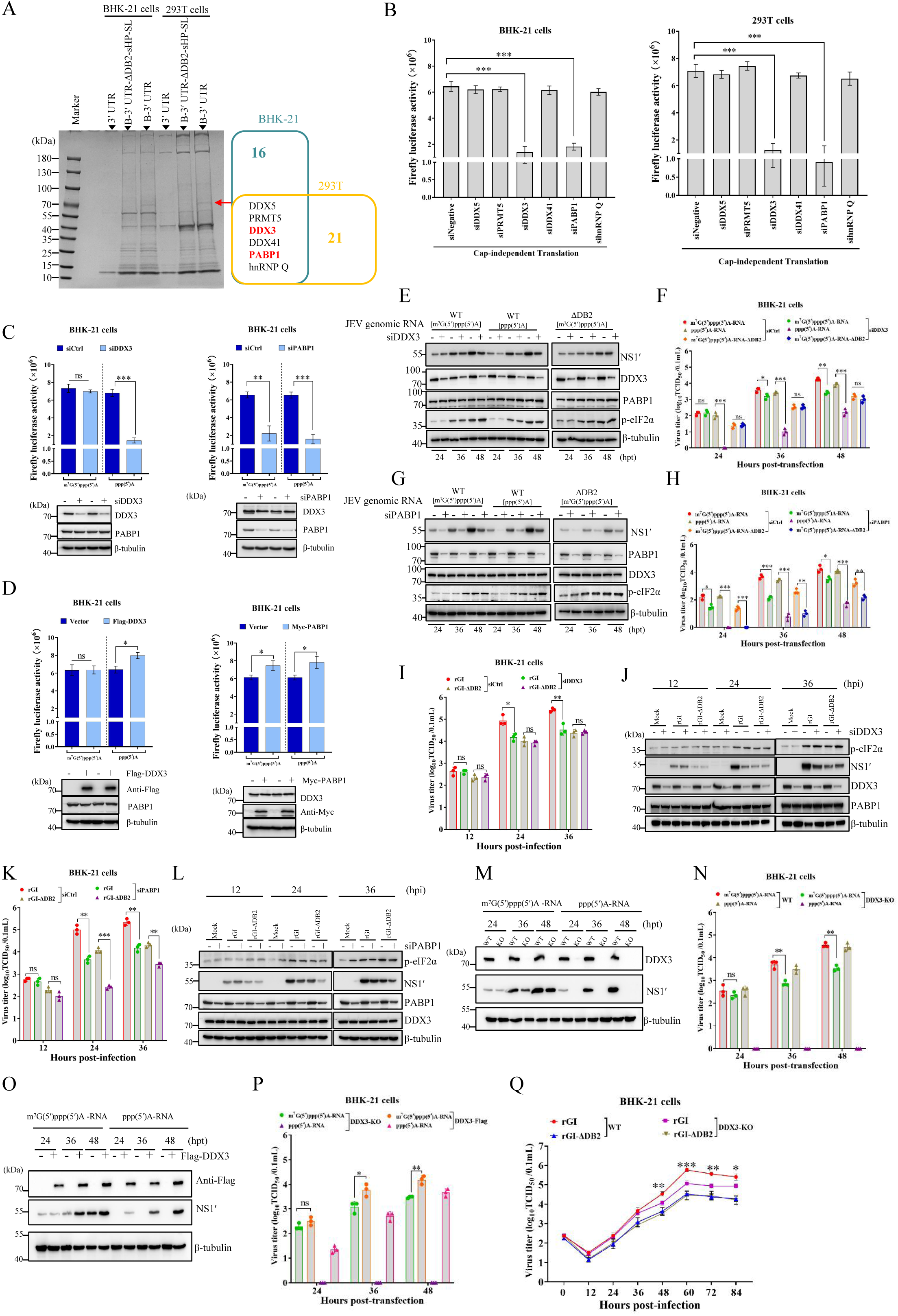
DDX3 and PABP1 regulate cap-independent translation initiation of JEV *in vitro*. (A) Identification of cellular proteins associated with JEV 3′UTR and JEV 3′UTR-ΔDB2-sHP-SL (left). Venn diagram of DB2 and sHP-SL specific binding proteins in BHK-21 and 293T cells detected by RNA pull-down and mass spectrometry (right). (B) The *firefly* luciferase activity assay using the noncapped monocistronic RNA reporter JEV-5′UTR-FLuc-3′UTR in BHK-21 and 293T cells with an indicated protein silenced using siRNAs (n=3). (C) BHK-21 cells pre-treated with 100 pmol of a mixture of DDX3-specific siRNAs (left) or PABP1-specific siRNAs (right) were respectively transfected with monocistronic RNA reporters JEV 5′ UTR-FLuc-3′ UTR with 5’terminal modified by either m^7^G(5′)ppp(5′)A or ppp(5′), and then harvested at 12 hours post-transfection for analysis of *firefly* luciferase activities (n=4). The levels of endogenous DDX3 and PABP1 were detected by western blotting. (D) The *firefly* luciferase activity assay of monocistronic RNA reporter JEV-5′UTR-FLuc-3′UTR in BHK-21 cells with Flag-DDX3 (left) or Myc-PABP1 (right) over-expressed (n=4). (E-H) BHK-21 cells with DDX3 or PABP1 silenced were transfected with the capped JEV genomic RNA of WT or ΔDB2 mutant, or noncapped WT genomic RNA. At the indicated time points post-transfection, the expression of viral NS1′ protein, p-eIF2α, DDX3 and PABP1 in BHK-21 cells was analyzed by immunoblotting (E and G), and viral titers in culture supernatants were determined by TCID_50_ assays (F and H) (n=3). (I-L) BHK-21 cells with DDX3 or PABP1 silenced were infected with rGI and rGI-ΔDB2 at a dose of 0.05 MOI. Replication of rGI and rGI-ΔDB2 was monitored by TCID_50_ assays (I and K) (n=3). The expression of viral NS1′ protein, p-eIF2α, DDX3 and PABP1 in BHK-21 cells was analyzed by immunoblotting (J and L). (M-P) The translational activity of JEV genomic RNA in DDX3-KO BHK-21 cells. DDX3-KO cells with or without DDX3-Flag over-expression were transfected with JEV genomic RNA m^7^G(5′)ppp(5′)A-RNA, ppp(5′)A-RNA, respectively. At different time points post-transfection, immunoblotting analysis of viral NS1′ protein (M and O). Virus titers in culture supernatant were evaluated by TCID_50_ assays (N and P) (n=3). (Q) The replication ability of rGI and rGI-ΔDB2 in WT and DDX3-KO BHK-21 cells (n=3). The significant differences between WT and DDX3-KO BHK-21 cells are marked (*, p<0.05; **, p<0.01; **, p<0.001). *, p<0.05; **, p<0.01; ***, p<0.001; ns, no statistical differences. Data are presented as mean ± SD of three independent experiments (B, C, D, F, H, I, K, N, P and Q).

The function of DDX3 and PABP1 in JEV replication was further evaluated. The siRNA-mediated knockdown of DDX3 expression dramatically attenuated rGI replication by 10-fold to 100-fold at 24 and 36 hpi but did not affect the replication of rGI-ΔDB2 that is defective in cap-independent translation (Figure 5I and 5J). Besides downregulating the titers of rGI at 24 and 36 hpi, PABP1 knockdown also exhibited an inhibitory effect on the replication of rGI-ΔDB2(Figure 5K and 5L). Meanwhile, we generated a DDX3-KO cell strain by editing the DDX3 gene but failed to obtain PABP1-KO cells. The expression of DDX3 in DDX3-KO cells was completely knocked out by a deletion of five nucleotides in exon 1 (Figure 5-figure supplement 2A-C). The knockout of DDX3 did not affect cell proliferation (Figure 5-figure supplement 2D). Virus recovery of rGI was attempted using WT and DDX3-KO BHK-21 cells. Only the transfection of DDX3-KO cells with the noncapped genomic RNA failed to rescue viable viruses, and viral titers rescued with the capped RNA in DDX3-KO cells were decreased (Figure 5M and 5N). Consistently, ectopically expressed DDX3-Flag in DDX3-KO cells rescued rGI with noncapped RNA and increased virus titers rescued with capped RNA (Figure 5O and 5P). Based on viral growth curves, rGI growth in DDX3-KO cells was significantly alleviated since 48 hpi compared to that in WT cells, while no obvious difference was observed in virus titers of rGI-ΔDB2 between WT and DDX3-KO cells (Figure 5Q). Taken together, the loss of function and gain of function experiments demonstrated that DDX3 and PABP1 are host factors positively regulating cap-independent translation of JEV, and PABP1 is also important for cap-dependent translation.

### The RNA structures of UTRs bound by DDX3 and PABP1 determine the cap-independent translation of JEV

An RNA pull-down assay was conducted to verify the interactions among DDX3, PABP1, and JEV 3′UTR. DDX3 and PABP1 were pulled down by biotinylated 3′UTR but not by biotinylated 3′UTR-ΔDB2-sHP-SL (Figure 6A), suggesting that DB2 and sHP-SL bridge the interactions between DDX3/PABP1 and JEV 3′UTR. In a competition test, the interactions DDX3/PABP1 with JEV 3′UTR were significantly out-competed by nonbiotinylated JEV 3′UTR in a dose-dependent manner, but not by JEV 3′UTR-ΔDB2-sHP-SL (Figure 6B). To exclude the interference of other host factors, purified GST-DDX3 and GST-PABP1 were used for RNA pull-down assay. GST-DDX3 and GST-PABP1, but not GST, coprecipitated with biotin-labeled JEV 3′UTR *in vitro* (Figure 6C), indicating that DDX3 and PABP1 directly bind to JEV 3′UTR. Moreover, DDX3 was pulled down by biotinylated 3′UTR and 3′UTR-ΔsHP-SL, while PABP1 was pulled down by biotinylated 3′UTR and 3′UTR-ΔDB2 (Figure 6D). These results demonstrated two pairs of direct interactions, DDX3/DB2 and PABP1/sHP-SL. In addition, the interaction between DDX3 and PABP1 was confirmed using co-IP (Figure 6E), GST pull-down (Figure 6F and 6G), and confocal microscopy (Figure 6H). Taken together, the interactions (Figure 6I) among DDX3, PABP1, and JEV 3′UTR (DB2 and sHP-SL) could be essential for JEV cap-independent translation.

**Figure 6.**
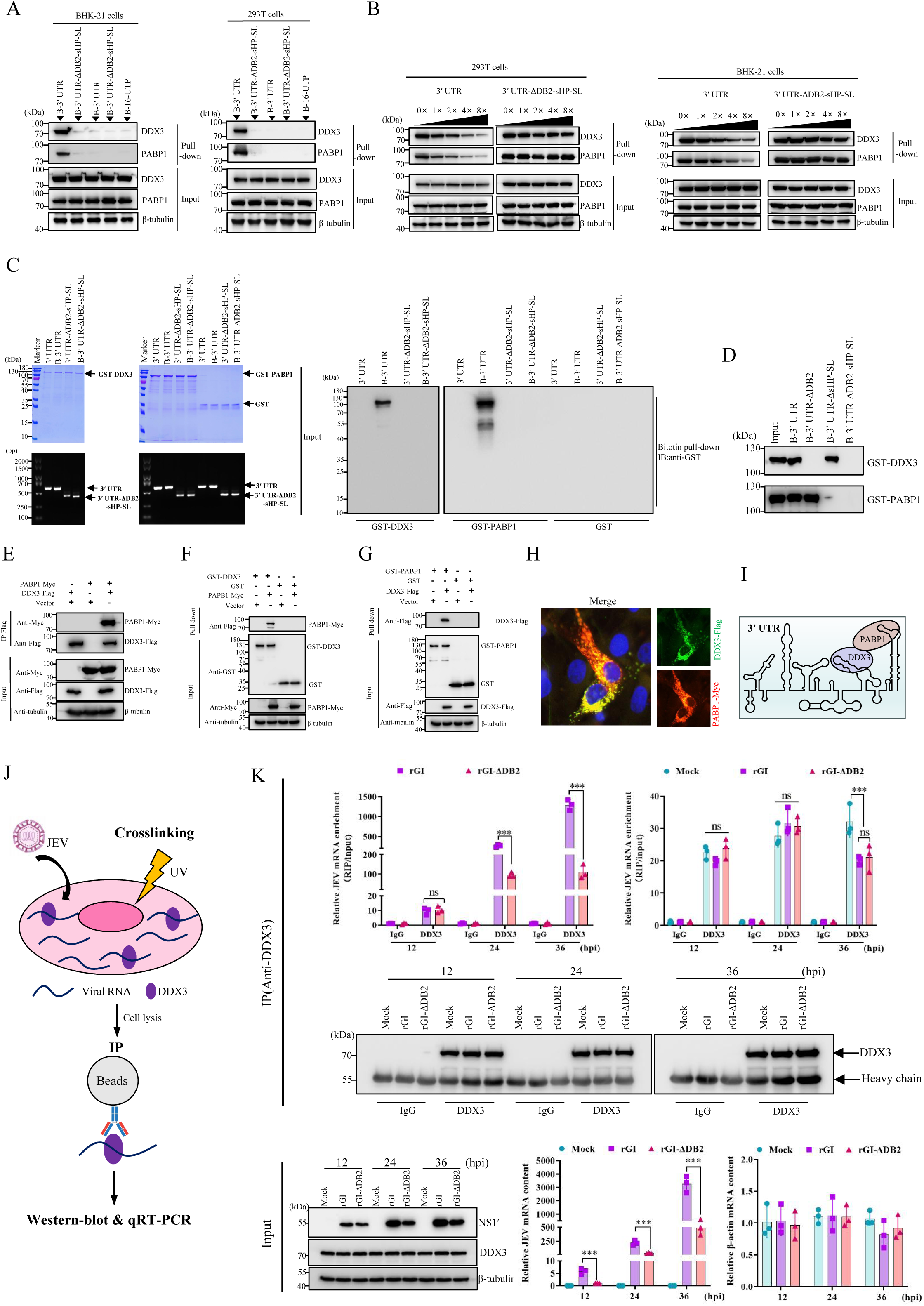
The interactions among DDX3, PABP1 and JEV 3′UTR. (A) The cell lysates of BHK-21 and 293T cells were respectively incubated with biotinylated-3′UTR (B-3′UTR), biotinylated-3′UTR-ΔDB2-sHP-SL (B-3′UTR-ΔDB2-sHP-SL), nonbiotinylated-3′UTR (3′UTR) or nonbiotinylated-3′UTR-ΔDB2-sHP-SL(3′UTR-ΔDB2-sHP-SL). The bound complexes were analyzed by immunoblotting. (B) The interactions between DDX3/PABP1 and JEV 3′UTR were confirmed by competition assays. (C and D) RNA pulldown analysis of the interaction between nonbiotinylated or biotinylated 3′UTR and recombinant proteins GST-DDX3, GST-PABP1, or GST. The bound complexes were analyzed by immunoblotting, and recombinant proteins and RNA in the input were detected by Coomassie brilliant blue staining and RT-PCR. (E) Co-immunoprecipitation analysis of the interaction between the ectopically expressed DDX3-Flag and Myc-PABP1 in HEK293T cells. (F and G) GST pulldown analysis of the interaction between recombinant DDX3 and PABP1. (H) Immunofluorescent analysis of the interaction between DDX3-Flag and PABP1-Myc in BHK-21 cells. (I) A model for the interactions among DDX3, PABP1, and JEV 3′UTR. (J) Schematic diagram of the RIP assay. BHK-21 cells are infected with rGI or rGI-ΔDB2 at an MOI of 0.01. At 12, 24 and 36 hours post-infection, the cells are cross-linked with short-wave UV light to form RNA-protein complexes, and then lysed for RNA immunoprecipitation. (K) Quantification of intracellular DDX3 protein expression via western blotting (Input, left), and JEV mRNA (Input, middle) and β-actin mRNA (Input, right) via RT-qPCR (n=3). Quantification of RIP-qPCR analysis illustrating the immunoprecipitation of JEV and β-actin mRNA upon incubation with either anti-IgG or anti-DDX3 beads (IP, n=3). The outcomes were calculated by normalization of RNA abundance in RIP samples (IgG and DDX3 mutant) to respective inputs (the analysis value of IgG group was set as 1). *, p<0.05; **, p<0.01; ***, p<0.001; ns, no statistical differences. Data are presented as mean±SD of three independent experiments

To examine the interaction between DDX3 and viral RNA in infected cells, we infected BHK-21 cells with either WT virus rGI or the mutant virus rGI-ΔDB2. At different time points post-infection, the cells were cross-linked with short-wave UV light to fix cellular RNA-protein complexes for RNA immunoprecipitation using a DDX3 antibody or an IgG control (Figure 6J). The enrichment of JEV RNA in the RNA-protein complex immunoprecipitated by the DDX3 antibody were analyzed by RT-qPCR. As shown in Figure 6K, in comparison with the IgG control, the DDX3 antibody pulled down significantly higher levels of JEV mRNA. In the RNA-protein complex immunoprecipitated by the DDX3 antibody, the levels of rGI- ΔDB2 genomic RNA were significantly lower than those of the WT genomic RNA at 24 hpi and 36 hpi. These results confirmed that DDX3 binds viral RNA in JEV-infected cells and DB2 of 3′UTR plays a key role in this interaction.

To pinpoint the key RNA structures in DB2 and sHP-SL mediated their interactions with DDX3 and PABP1, a panel of JEV 3′UTR mutants was generated by introducing mutations or deletions (Figure 7-figure supplement 1A). All mutants except ΔDB2 and DB2mutΔ1-4 bind to DDX3, and the binding of DDX3 to DB2mut1, DB2mut4, DB2mutΔ1, and DB2mutΔ5 was reduced (Figure 7A). Using the monocistronic reporters containing the corresponding mutations at DB2, the disrupted RNA structures in DB2 by DB2mut1 and DB2mut4 consistently decreased the cap-independent FLuc expression (Figure 7B). Meanwhile, the capped and noncapped viral genomic RNAs with the corresponding DB2 mutations or deletions were used to rescue viruses. Recombinant viruses were successfully recovered using all RNA transcripts except for noncapped-ΔDB2, DB2 mutΔ1 and DB2mutΔ1-4 which are unable to bind to DDX3 (Figure 7E, Table supplement 2). Thus, the disrupted RNA structures in DB2 by DB2mut1 and DB2mut4 mediate the interaction between DB2 and DDX3, which is also essential for the cap-independent translation of JEV. Meanwhile, using the JEV 3′UTR mutants with modified sHP-SL (Figure 7-figure supplement 1B), the loop1 and loop2 structures of SL disrupted by mut4 and SL mutΔ2-3 was identified as the key region that mediated the interaction between 3′UTR and PABP1 (Figure 7C). Consistently, this key structure is also critical for the cap-independent FLuc expression in the reporter assay (Figure 7D). The corresponding mutations or deletions in sHP-SL were also introduced into the viral genome to rescue viruses. We did not detect the viral NS1′ protein expression and failed to rescue viruses with ΔsHP-SL and SLmut4 mutants (Figure 7F, Table supplement 2). Of note, unlike the ΔsHP-SL mutant, the SLmutΔ2-3 mutant was rescued using capped viral genomic RNA but not the noncapped one (Table supplement 2). Therefore, we eventually identified the key RNA structures in DB2 and sHP-SL of 3′UTR which regulate JEV cap-independent translation through their interactions with cellular DDX3 and PABP1.

**Figure 7.**
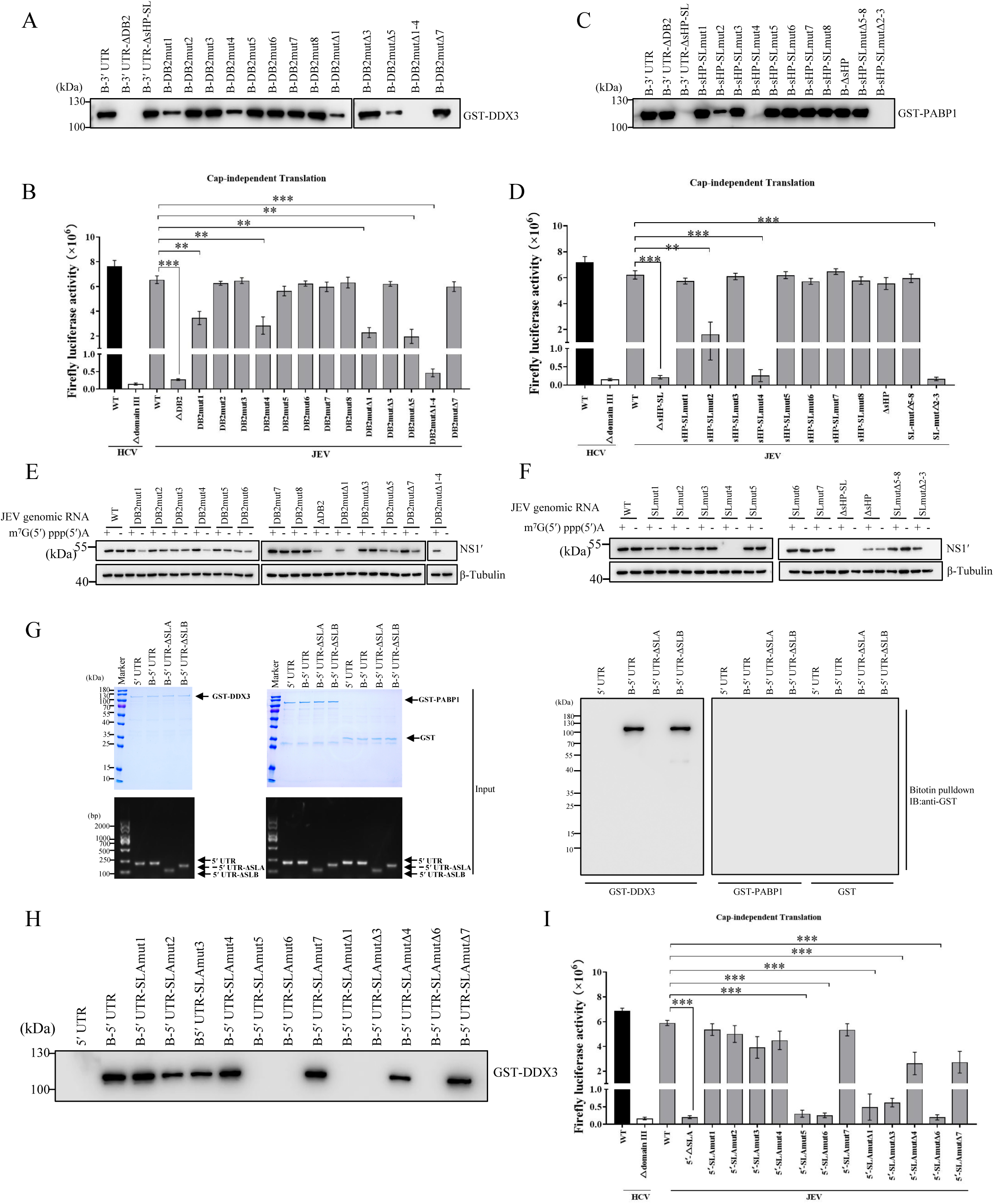
The key RNA structures in UTRs bound by DDX3 and PABP1 are critical to the cap-dependent translation initiation of JEV. (A and C) Mapping interaction regions in elements DB2 and sHP-SL for DDX3 and PABP1. Pulldown assays were performed with biotinylated-JEV 3′UTR or 3′UTR mutants and the purified recombinant GST-DDX3(A) or GST-PABP1 (C). The bound complexes were analyzed by immunoblotting. (B, D and I) The *firefly* luciferase activity assay of JEV-RLuc reporter mutants in BHK-21 cells. BHK-21 cells were transfected with the indicated noncapped reporters, and then lysed at 12 hours post-transfection for analyzing the luciferase activities (n=4). (E and F) Western-blot analysis of BHK-21 cells transfected with the capped and noncapped viral genomic RNAs carrying the corresponding DB2 and sHP-SL mutations or deletions. (G) RNA pulldown analysis of the interaction between nonbiotinylated or biotinylated 5′UTR and recombinant GST-DDX3, GST-PABP1, or GST. The bound complexes were analyzed by immunoblotting, and recombinant proteins and RNA in the input were detected by Coomassie brilliant blue staining and RT-PCR. (H) Mapping regions in 5′UTR bound by DDX3. **, p < 0.01; ***, p < 0.001. Data are presented as mean ± SD from four independent experiments (B, D and I).

Previous studies have revealed that DDX3 binds to both UTRs of the JEV genome to regulate viral translation(Li et al., 2014). We further explored whether DDX3 or PABP1 binds to 5′UTR to regulate JEV cap-independent translation. In line with a previous study (Li et al., 2014), DDX3, instead of PABP1, directly bind to the SLA structure of 5′UTR in the RNA pull-down assay (Figure 7G). A panel of 5′UTR mutants was generated to narrow down the key RNA structures in SLA mediating the interaction between 5′UTR and DDX3 (Figure 7-figure supplement 1B). The GST pull-down results revealed that DDX3 interacted directly with the stem-loop1 and stem-loop2 structures of SLA because the modifications of this structure reduced or diminished the interaction (Figure 7H). In the cap-independent translation assay, the 5′UTR mutants that can’t interact with DDX3 are defective in FLuc expression (Figure 7I), confirming the involvement of 5′UTR in JEV cap-independent translation through its interaction with DDX3.

### DDX3 does not rely on its RNA helicase activity to regulate JEV cap-independent translation

Since DDX3 belongs to the DEAD box RNA helicase family that harbor ATPase and RNA helicase activities(Winnard et al., 2021), we sought to determine whether RNA helicase activity of DDX3 is required for JEV cap-independent translation. The DDX3-Mut containing mutations at the ATPase motif (K to E) and helicase motif (S to L) of DDX3 was constructed to eliminate its RNA helicase activity as described previously (Wang et al., 2009; Yedavalli et al., 2004) (Figure 8A). The recovery of rGI was attempted using DDX3-KO cells with or without the overexpression of DDX3. Recombinant rGI was rescued using noncapped RNA transcript with the help of DDX3-WT-Flag or DDX3-Mut-Flag, while virus recovery failed in DDX3-KO cells (Figure 8B and 8C). Meanwhile, the downregulation of DDX3 helicase activity by the specific inhibitor RK33(Xie et al., 2016) in BHK-21 cells, had no obvious effect on NS1′ expression and viral yields (Figure 8D and 8E), suggesting that DDX3 does not rely on its RNA helicase activity to regulate JEV cap-independent translation.

**Figure 8.**
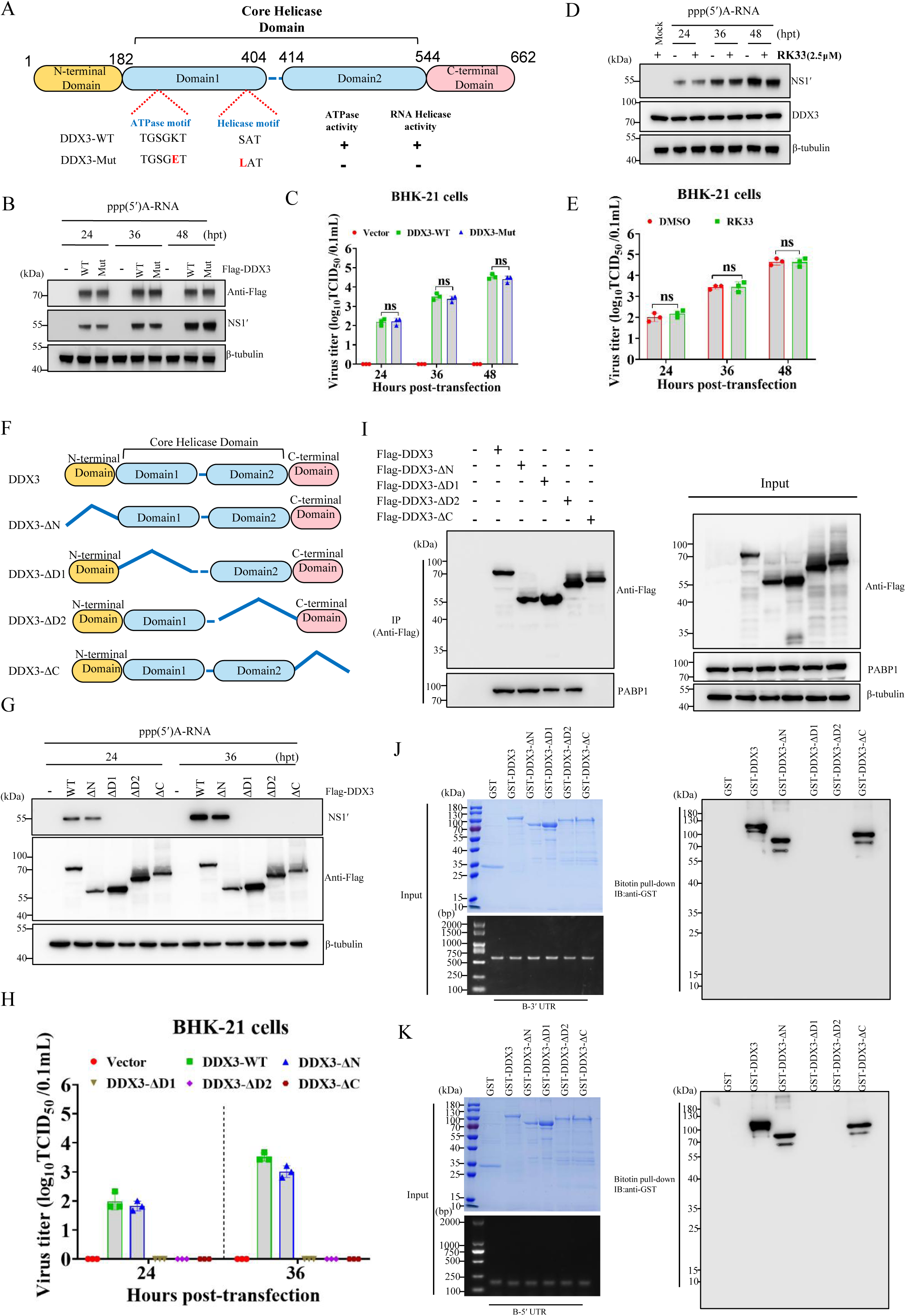
The regulation of JEV cap-independent translation by DDX3 relys on its interaction with JEV UTRs and PABP1 but not its helicase activity. (A) Schematic representation of the WT and mutant DDX3. The lysine (K) in the ATPase motif and the serine (S) in the helicase motif were respectively changed into glutamic acid (E) and leucine (L) in DDX3-Mut. +, presence of ATPase or RNA helicase activities; -, absence of ATPase or RNA helicase activities. (B and C) The translational activity of JEV genomic RNA in DDX3-KO BHK-21 cells. DDX3-KO cells with DDX3-WT-Flag or DDX3-Mut-Flag overexpression were transfected with noncapped JEV genomic RNA. At different time points post-transfection, immunoblotting analysis of viral NS1′ protein (B). Virus titers in culture supernatants were evaluated by TCID_50_ assays (C) (n=3). (D and E) BHK-21 cells were treated with RK-33 (2.5μM), and then transfected with noncapped JEV genomic RNA of WT. At indicated time points post-transfection, NS1′ protein was detected by immunoblotting (D) and viral titers in culture supernatants were determined by TCID_50_ assays (E) (n=3). (F) Schematic representation of the truncated mutants of DDX3. (G and H) The translational activity of JEV genomic RNA in DDX3-KO BHK-21 cells with the overexpression of DDX3 or DDX3 mutants. (I) Co-immunoprecipitation analysis of the interaction between the ectopically expressed DDX3-Flag or its mutants and Myc-PABP1 in HEK-293T cells. (J and K) RNA pulldown analysis of the interaction between biotinylated 5′UTR or 3′UTR and recombinant GST-DDX3, GST-DDX3-ΔN, GST-DDX3-ΔD1, GST-DDX3-ΔD2, GST-DDX3-ΔC or GST. Data are presented as mean ± SD of three independent experiments (C, E and H). **, p<0.01; ***, p<0.001; ns, no statistical difference.

DDX3 contains a highly variable N-terminal domain core-helicase domain, a core-helicase domain that is composed of two Rec-like domains (D1/2) in tandem and a C-terminal domain (Figure 8A). To determine which domain of DDX3 is essential for JEV cap-independent translation, we constructed several DDX3 mutants with deletions and assessed their function in DDX3-KO cells (Figure 8F). We found that the deletion of any domains other than the N-terminal domain led to the loss of viral protein expression and the failure of virus recovery using noncapped viral RNA transcript in DDX3-KO cells (Figure 8G and 8H). Furthermore, the interaction between DDX3 and PABP1 was disrupted by the C-terminal domain deletion of DDX3 (Figure 8I), and the absence of either helicase domains resulted in the loss of the ability of DDX3 to bind to the 5′ and 3′UTRs of JEV (Figure 8J-8K). Thus, the C-terminal domain and helicase domains of DDX3 respectively mediate its interactions with PABP1 and JEV UTRs, which are critical for the cap-independent translation of JEV.

### DDX3 interacts with PABP1 to form a translation initiation complex for recruiting 43S PIC to JEV 5′UTR

We next attempted to gain insight into the mechanisms of DDX3 and PABP1 cooperatively regulating the cap-independent translation initiation of JEV. DDX3 and PABP1 have been reported to interact with the 40S ribosomal subunit to support the assembly of functional 80S ribosomes in eukaryotic cells(Geissler et al., 2012; Soto-Rifo et al., 2012). The distribution of DDX3 and PABP1 in separate ribosomal subunits (40S and 60S), monosomes (80S), and larger polysomes was evaluated by polysome profiling. Based on the absorptions at 254 nm, the obtained fractions showed a distinguishable profile with peaks representing four ribosome states (Figure 9A). Ribosomal protein RPS6 in the fractions containing ribosomes was detected by Western blot (Figure 9B). In cells transfected with uncapped JEV genome of ΔDB2-sHP-SL mutant or mock-transfected, DDX3 mainly distributed in the top fractions without ribosomes but not in the larger polysomes fractions, while most of PABP1 sedimented in the 40S and 60S fractions and partially tailed into the 80S fractions (Figure 9B). Of note, the distribution of DDX3 shifted significantly to the fractions of 40S, 60S and 80S ribosome states in cells transfected with noncapped JEV genome. These data supported that DDX3 specifically participates in the translation initiation phase of the noncapped JEV genome, and PABP1 is involved in the initiation phase of canonical translation.

**Figure 9.**
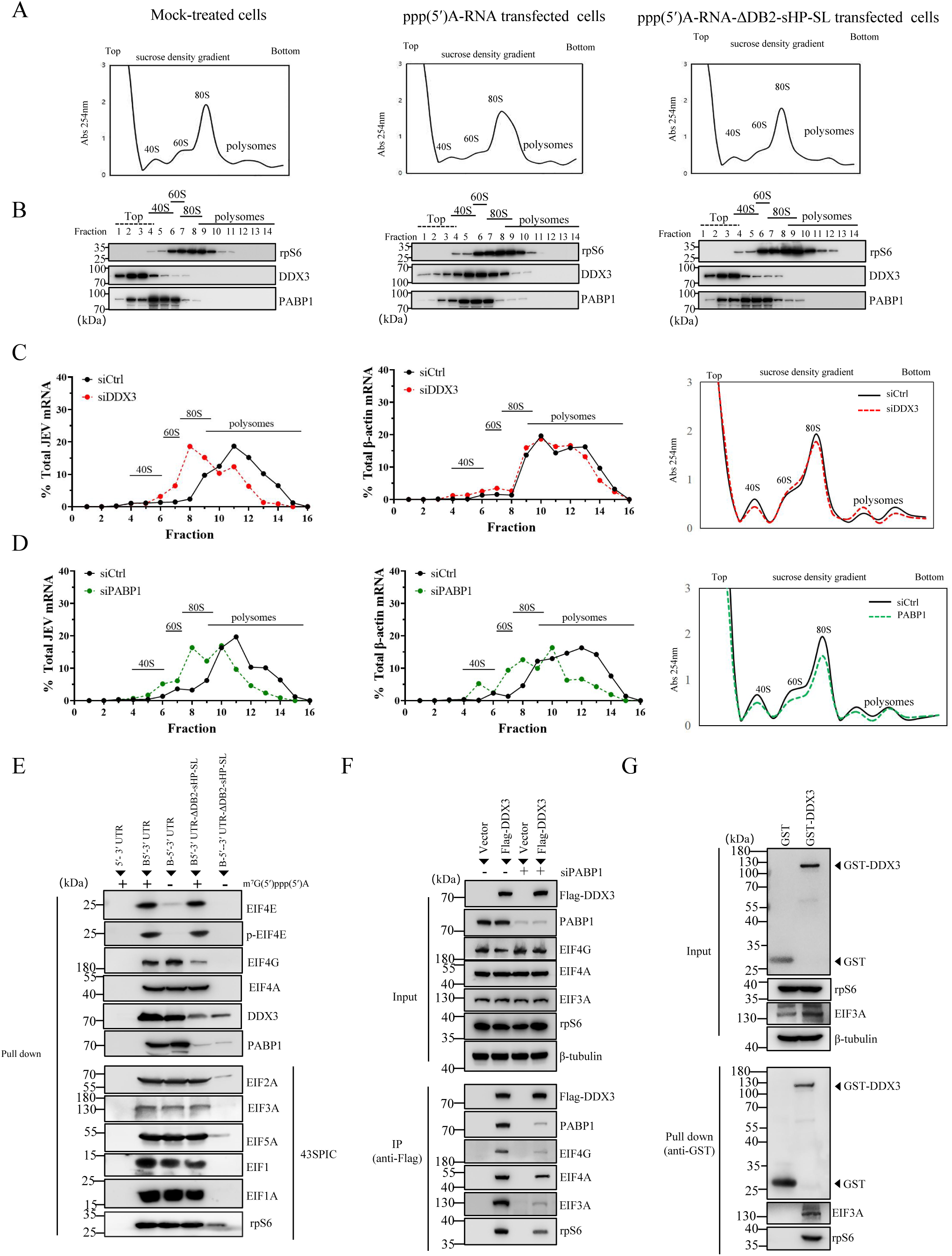
DDX3 cooperates with PABP1 to recruit translation initiation factors and 43S PIC for optimal cap-independent translation initiation of JEV. (A and B) Ribosome profiles of BHK-21 cells mock-transfected or transfected with noncapped JEV genome or ΔDB2-sHP-SL mutant. The ribosome profiles were obtained by measuring the absorbance at 254 nm of individual fractions(A). Each fraction was immunoblotted using the antibodies rPS6, DDX3 and PABP1 (B). (C and D) BHK-21 cells treated with either a control siRNA (siCtrl) or siRNA targeting DDX3 (C) or PABP1 (D) were transfected with JEV ppp(5 ′) A-RNA and ppp(5′)A-RNA-ΔDB2-sHP-SL at a dose of 2μg. At 8 hours post-transfection, cell lysates were resolved and fractionated through sucrose gradients. Each fraction was subjected to RT-qPCR analysis for JEV and β-actin mRNA levels. The percentage of mRNA transcripts recovered from each fraction was plotted against the fraction number. (E) RNA pulldown analysis of host factors bound by nonbiotinylated or biotinylated JEV reporter RNA. The input and the precipitates were analyzed by western blotting with the indicated antibodies. (F) BHK-21 cells with or without the treatment of siPABP1 were transfected with Flag-DDX3 for 24 hours and then lysed for immunoprecipitation with an anti-Flag antibody. The protein complex was analyzed by immunoblotting. (G) GST pulldown analysis of interacting partners of recombinant GST and GST-DDX3 in BHK-21 cell lysate.

Whether DDX3 and PABP1 function differently in cap-dependent and cap-independent translation was further investigated. The distribution profiles of noncapped JEV mRNA and β-actin mRNA were determined by their percentages in the fractions of BHK-21 extracts (Figure 9C and 9D). The distribution of JEV mRNA and β-actin mRNA gradually increased from monosome to polysome with peaks in fractions 10 to 12 in control cells, suggesting effective ribosome loading and continuous translation. The distribution of JEV mRNA shifted considerably toward lighter fractions with a peak at fractions 7 to 9 when DDX3 was knocked down, while the distribution of β-actin mRNA was not affected (Figure 9C), suggesting that DDX3 is a host factor specifically required for cap-independent translation but not for canonical translation. In contrast, the distribution of JEV mRNA and β-actin mRNA shifted considerably toward lighter fractions when PABP1 was depleted (Figure 9D), confirming that PABP1 is critical for both cap-dependent and cap-independent translation mechanisms.

To further clarify the function of DDX3 at the initiation step of JEV cap-independent translation, affinity purification with the capped and noncapped biotinylated RNAs was performed to analyze translation initiation factors recruited by DDX3 to JEV 5′UTR. A range of host factors associated with translation initiation was recruited to the noncapped JEV-UTRs, including eIF4G and eIF4A, as well as the components of 43S PIC (Figure 9E). As expected, EIF4E and p-EIF4E were recruited to the capped RNAs but not the noncapped RNAs. In the absence of DB2 and sHP-SL, dramatically reduced DDX3 and PABP1 were pulled down by biotinylated RNA of JEV-UTRs, which led to the loss of other factors associated with translation initiation using the noncapped RNA. The recruitment of translation initiation complexes by DDX3 was further evaluated by co-IP assay. DDX3 specifically coprecipitated with PABP1, eIF4G, and eIF4A, as well as eIF3A and rpS6 in the 43S complex, and PABP1 depletion alleviated these interactions (Figure 9F). The interactions between DDX3 and eIF3A/rpS6 were confirmed by the GST pull-down assay (Figure 9G). In addition, PABP1 recruiting eIF4G to initiate translation has been demonstrated(Kahvejian et al., 2001; Ma et al., 2009), and the interaction between DDX3 and PABP1 was confirmed in Figure 6E-H. Thus, our results suggested that DDX3 could interact with PABP1 to form DDX3/PABP1/eIF4G/eIF4A tetrameric complex, thereby recruiting 43S PIC to the 5′ end of JEV genome and allowing cap-independent translation initiation.

## DISCUSSION

As obligate intracellular parasites, viruses exclusively rely on the host translation apparatus to support the synthesis of viral proteins. Upon infection, viruses employ diverse mechanisms to prevent the efficient expression of cellular proteins, while viral protein expression could evade these inhibitory mechanisms(Walsh et al., 2013; Walsh and Mohr, 2011). During flavivirus infection, the global cellular protein synthesis is almost completely repressed by PKR-induced phosphorylation of eIF2α (Tu et al., 2012; Wang et al., 2020), the inhibition of rRNA synthesis(Selinger et al., 2019), or other diverse mechanisms(Roth et al., 2017). Contrary to cellular mRNA, viral mRNA is usually translated efficiently thanks to alternative translation initiation strategies. Here, we revealed that JEV translation remains efficient when the cap-dependent translation was dramatically inhibited in mammalian, avian, and mosquito cells, suggesting a possible switch of JEV translation initiation from cap-dependent to cap-independent under host translational shutoff. This prompted us to investigate the mechanism of the selective synthesis of viral protein in a cap-independent manner during JEV infection.

Two alternative but not mutually exclusive cap-independent mechanisms of translation initiation have been proposed in flaviviruses. One mechanism is IRES-dependent initiation(Fernández-García et al., 2021; Song et al., 2019), while the other mechanism is dependent on the 3′-end genome(Berzal-Herranz et al., 2022; Edgil et al., 2006; Wang et al., 2020). To date, there is no consensus regarding whether the cap-independent translation of flaviviruses is IRES-dependent or not. Like the other flaviviruses(Song et al., 2019), JEV can be rescued with the noncapped genomic RNA in mammalian, avian, and insect cells, confirming the cap-independent translation of JEV. In a bicistronic reporter assay for IRES identification, the 5′UTR or 5′UTR-cHP-cCS of the JEV genome enabled extremely low levels of cap-independent translation. This is consistent with a previous reported “IRES-like” activities of the flavivirus 5′UTRs resulting in less than 1% of the EMCV IRES activity(Edgil et al., 2006; Wang et al., 2020). The typical IRES is approximately 300 nucleotides in length and is characterized by complicated high-order RNA structures assembled with stem-loops and pseudoknots which directly recruit translation initiation factors eIF4G/4A and the 43S PIC(Filbin and Kieft, 2009; Lee et al., 2017). In contrast, the length of flavivirus 5′UTR is always less than 110 nucleotides, and JEV 5’UTR can’t recruit eIF4G/4A and 43S PICs. Thus, it is reasonable that the 5′UTR of JEV as well as other flaviviruses does not harbor IRES activity. Our results, as well as previous data(Alvarez et al., 2005; Edgil et al., 2006; Wang et al., 2020), only demonstrated flavivirus 5′UTR does not have IRES-like activity. However, we can’t rule out the possibility that flavivirus 5′UTR together with some following coding sequences could form an IRES-like structure.

Viral translation could be enhanced by the 3′UTR of their genomes (Mazumder et al., 2003; Rasekhian et al., 2021). In flavivirus, in addition to its role in viral RNA replication and virus production, 3′UTR serves as a translation initiation enhancer which functions similarly to the 3′-poly(A) tail in enhancing cap-dependent translation(Berzal-Herranz et al., 2022; Holden and Harris, 2004; Ramos-Lorente et al., 2024). Intriguingly, the 3′UTR of flaviviruses can equally stimulate cap-independent translation in the presence of flavivirus 5′UTR as well as other cellular and viral 5′UTRs. For instance, the 3′UTR of DENV stimulated the translation of reporter mRNAs containing the noncapped 5′UTRs of EMCV, BVDV, or human β-globin(Edgil et al., 2006; Holden and Harris, 2004), and the 3′UTRs of WNV, DENV, and YFV promoted the cap-independent translation driven by WNV 5′UTR (Berzal-Herranz et al., 2022). In line with these reports, in the presence of 3′UTR, JEV 5′UTR enabled the cap-independent translation. In addition, the cap-independent translation mediated by JEV 5′UTR remained efficient when JEV 3′ UTR was substituted with the 3′UTRs of other flaviviruses, further supporting that the 3′UTR is essential for the cap-independent translation of flaviviruses. In all flaviviruses, an array of RNA secondary structures has been predicted with their 3′UTRs which are about 400 to 800 nucleotides in length, including highly complex RNA structures that are closely related to viral translation efficiency, viral replication, and pathogenicity of flaviviruses (Upstone et al., 2023; Villordo et al., 2016; Zhang et al., 2022). In this study, DB2 and sHP-SL in JEV 3′UTR were identified as key RNA structures for the cap-independent translation of JEV. Furthermore, the recovery of rGI-ΔDB2 with the capped viral genome but not the noncapped one, confirms the decisive role of DB2 in the cap-independent translation of JEV. Previous studies suggested that the 3′SL of sHP-SL enhances translation by increasing mRNAs recruited into polysomes and the number of ribosomes associated with mRNA, thereby increasing translation initiation(Holden and Harris, 2004; Ramos-Lorente et al., 2024). Here, we also determined that sHL-SL contributed to maximizing cap-dependent translation efficiency mediated by JEV 5′UTR which is indispensable for the cap-independent translation. The deletion of DB2 resulted in decreased replication capacity of JEV in BHK-21 cells and reduced JEV virulence in mice, while the deletion of sHP-SL is lethal for viral viability. These results suggested that the cap-independent translation of JEV is not only important for its evasion of host shutoff *in vitro* but also for its virulence *in vivo*.

A closed-loop architecture is required for efficient translation of most mRNAs(Kahvejian et al., 2001; Thompson and Gilbert, 2017). Similarly, the circular topology of the flaviviral RNA genome during translation has also been proposed(Chiu et al., 2005; Kofler et al., 2006). A defining feature of flavivirus genomes is the existence of complementary sequence motifs at both termini that are involved in the genome circularization(Alvarez et al., 2005). JEV 5′UTR can’t support cap-independent translation without JEV 3′UTR, and JEV 3′UTR in stimulating cap-independent translation was not affected by the absence of the cyclization sequence (CS) of 5′UTR. These observations suggested the cap-independent translation initiation of JEV may not require direct interaction between 5′UTR and 3′ UTR. We hypothesized that host factors recruited by JEV UTRs are required for the cap-independent translation initiation. Therefore, a panel of cellular proteins interacting with DB2 and sHP-SL of 3′UTR was identified. Among them, DDX3 and PABP1 were demonstrated to regulate JEV cap-independent translation. Both DDX3 and PABP1 were considered as the general, auxiliary translation factors involved in the translation initiation process(Geissler et al., 2012; Lee et al., 2008; Wang et al., 2022). PABP1 bound to the poly (A) tail interacts with the translation initiation complex eIF4F bound to the cap structure of mature mRNAs, thus contributing to the formation of a closed loop structure that enhances translation or promotes translation re-initiation(Kahvejian et al., 2001). In flavivirus infection, PABP1 could also serve as a bridging factor binding 3′UTR internally for circularization of the viral genome and translation enhancement(Polacek et al., 2009; Shen et al., 2022). In this study, PABP1 depletion reduced the efficiency of both cap-dependent and cap-independent translation of JEV, supporting that PABP1 exerts a regulatory function on the translation of flaviviruses. In contrast, DDX3 is specifically required for the cap-independent translation of JEV. As reported previously(Li et al., 2014), DDX3 binds to both UTRs of JEV mediated by the RNA structures of loop1-stem-loop2 in DB2 and loop1-stem1/2-loop2 in SLA. In line with these results, DDX3 has been proven to be required for translation driven by cellular and viral transcripts that contain complicated RNA structures in their 5′UTR but not for general translation(Ayalew et al., 2016; Lai et al., 2008). Thus, DDX3 binds to UTRs to form a closed-loop architecture of the JEV genome.

DDX3 is a cellular ATP-dependent RNA helicase involved in different aspects of RNA function ranging from transcription to translation(Mo et al., 2021). An important aspect of DDX3 function is involved in translation, specifically at the initiation step(Lai et al., 2008). Of note, DDX3 appears to promote the translation of a subset of specific mRNA carrying highly structured 5′UTRs through its helicase activity which facilitates translation through the resolution of secondary structures during ribosomal scanning(Lai et al., 2010; Lai et al., 2008; Soto-Rifo et al., 2012). In this study, the knockout of DDX3’s helicase activity by site-directed mutagenesis or using the specific inhibitor RK33 had no obvious effect on NS1′ expression and viral yields of noncapped JEV genome, suggesting that the RNA helicase activity of DDX3 is important for its regulatory function in JEV cap-independent translation. Some functions of DDX3 depend on its helicase activity, while the others depend only on its direct interaction with RNA or proteins (Hernández-Díaz et al., 2021; Putnam and Jankowsky, 2013). In this study, we found that the C-terminal domain and helicase domains of DDX3 respectively mediate its interactions with PABP1 and JEV UTRs. The deletion of these domains of DDX3 led to a loss of cap-independent translation with noncapped JEV genome indicated by the failure of viral protein expression and virus recovery. Thus, the function of DDX3 in regulating JEV cap-independent translation depends only on its direct interactions with JEV UTRs and PABP1 but not its helicase activity.

In the translation initiation phase, DDX3 could remodel the RNA structures and remove the RNA-bound proteins of 5′UTR to facilitate 43S ribosome binding(Calviello et al., 2021; Soto-Rifo et al., 2012). In this study, we observed higher levels of DDX3 distributed in the fractions of ribosomal subunits and monosomes in cells transfected with noncapped JEV genome, in comparison to cells without JEV cap-independent translation. Thus, DDX3 may be actively involved in cap-independent initiation by assisting the assembly of functional 80S ribosomes for optimal translation, which was supported by the decreased distribution of JEV mRNA in the polysome fractions of DDX3 knockdown cells. Circularization of cellular mRNAs mediated by the PABP/eIF4G binary complex brings together the cap-binding protein eIF4E and the RNA helicase eIF4A (Kahvejian et al., 2001; Wells et al., 1998). Translation initiation is promoted by the interaction between eIF4G and eIF3 of the 43S PIC(Kahvejian et al., 2001). The noncapped JEV-UTRs specifically recruited PABP1, eIF4G, and eIF4A, as well as the components of the 43S PIC, which was dramatically inhibited by the deletion of DB2 and sHP-SL in 3′UTR. In addition, DDX3 coprecipitated with eIF4G, eIF4A, PABP1 and the components of the 43S PIC. These findings support that DDX3 recruits translation initiation factors eIF4G and eIF4A to JEV UTRs via its interaction with PABP1, thereby recruiting the 43S PIC to the 5′ end of the JEV genome and allowing the initiation of cap-independent translation.

In summary, we found that JEV employs a cap-independent translation strategy to evade host translational shutoff. JEV 5′UTR lacks IRES or IRES-like activity, while 5′UTR can initiate cap-independent translation in the presence of 3′ UTR. DB2 and sHP-SL of 3′UTR were involved in the regulation of cap-independent translation. DDX3 and PABP1 that interact with DB2 and sHP-SL are indispensable for JEV cap-independent translation. DDX3 binds to the loop1-stem1-loop2 structure of DB2 in 3′UTR and the stem-loop1 and stem-loop2 structures of SLA in 5′UTR, while PABP1 binds to the loop1-loop2 structure of sHP-SL in 3′UTR. Mechanistically, DDX3 binds to JEV UTRs to establish a closed-loop architecture, and its interaction with PABP1 to form DDX3/PABP1/eIF4G/eIF4A tetrameric complex, thereby recruiting the 43S PIC to initiate translation (Figure 10). Our findings demonstrated the cap-independent translation mechanism of JEV which relies on the High-order RNA structures formed by UTRs and the regulatory roles of DDX3 and PABP1. These results provide new insights into the understanding of flavivirus translation regulation.

**Figure 10.**
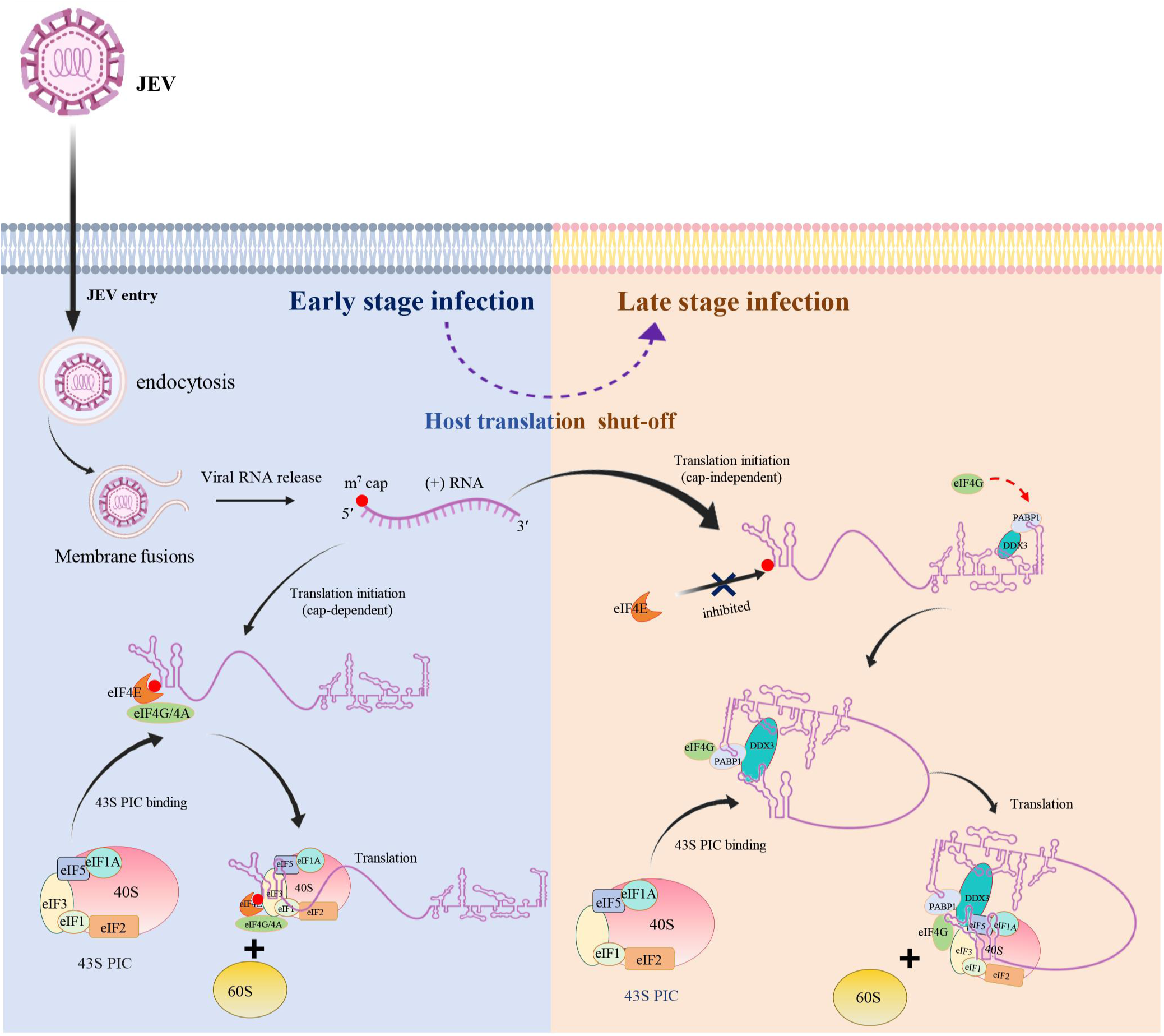
Schematic model of DDX3 and PABP1 regulation of JEV cap-independent translation initiation. During the shut-off of host cellular canonical translation, DDX3 and PABP1 respectively bind to *cis*-acting elements DB2 and sHP-SL, in which DDX3 could also anchor to 5′UTR of JEV genomic RNA to establish a closed-loop architecture. Simultaneously, DDX3 interacts with PABP1 to form DDX3/PABP1/eIF4G/eIF4A tetrameric complex, thereby recruiting 43S PIC to the 5’end of the viral genome and allowing translation initiation.

## Limitations of the study

Flaviviruses, like JEV, Zika virus, and TMUV, share some highly conserved RNA structures in 3′UTR. The reporter assay suggested that the conserved DB2 and sHP-SL structures in their 3′UTRs are crucial for the cap-independent translation of these viruses. Based on these observations, we speculated that the cap-independent translation of flaviviruses may be highly conserved. However, there are many other members in the genus of *orthoflavivirus* that have not been included in this study. To solid confirm this hypothesis, more experiments using other flaviviruses are required. In addition, due to the essential role of PABP1 in cellular translation, the PABP1-KO cells were not generated through CRISPR/cas9 gene editing to confirm its function in bridging the interactions between DDX3 and translation initiation factors in cap-independent translation.

## Materials and methods

### Reagents and antibodies

Mouse monoclonal anti-Flag antibody (M185-3L-1) was purchased from MBL. Mouse monoclonal anti-NS1′antibody (GTX633961) was purchased from GeneTex. Mouse monoclonal anti-Myc (M4439) and mouse anti-puromycin (MABE343) antibodies were purchased from Merck Millipore. Mouse anti-DDX3 (67915-1-Ig), mouse anti-PABP1 (66809-1-Ig), mouse anti-eIF4G (67199-1-Ig), mouse anti-rPS6 (66886-1-Ig), rabbit anti-eIF5 (11155-1-AP), rabbit anti-eIF1(15276-1-AP) antibodies were purchased from Proteintech Group. Rabbit anti-β tubulin(AP004) and rabbit anti-Flag antibodies(AP007) were purchased from Bioworld Technology. Mouse anti-GST (AE001), rabbit anti-Myc (AE009), rabbit anti-eIF4E (A2162), rabbit anti-Phospho-eIF4E (AP0747), rabbit anti-Phospho-eIF2α (A0764), rabbit anti-eIF4A (A24293), rabbit anti-eIF2A (A9709), rabbit anti-eIF3A(A0573), rabbit anti-eIF1A (A20954) antibodies were purchased from ABclonal Technology. HRP-conjugated goat anti-mouse (D11087) and HRP-conjugated goat anti-anti-rabbit (D11058) antibodies were purchased from BBI Life Science. Alexa Fluor 488-AffiniPure Goat Anti-Mouse IgG(H+L) Jackson (115-545-003) and TRITC-conjugated anti-rabbit (115-025-003) antibodies were purchased from Jackson ImmunoResearch. The chemical reagents puromycin (58-58-2) was purchased from Merck Millipore. The inhibitor 4E2RCat (HY-100733) and RK-33(HY-100455) were purchased from MedChemExpress.

### Cell lines

BHK-21, ST, DF-1, and HEK-293T cells were purchased from American Type Culture Collection (ATCC) and cultured in Dulbecco modified Eagle medium (DMEM, Hyclone) containing 10% fetal bovine serum (FBS, Sigma), 100 μg/mL streptomycin and 100 IU/mL penicillin (Hyclone) at 37°C in a 5% CO_2_ incubator. C6/36 cells purchased from ATCC were grown in RPMI medium 1640 supplemented with 10% FBS, 100 μg/ml streptomycin and 100 IU/ml penicillin at 28°C with no additional CO_2_. For the following viral infection, all cells were maintained in a medium containing 2% FBS, 100 μg/mL streptomycin and 100IU/mL penicillin.

### Ethics Statement and Biosafety Procedures

All animal experiments were approved by the Institutional Animal Care and Use Committee of Yangzhou University (IACUC No: SYXK-SU-202202163) and were performed in compliance with the Guidelines on the Humane Treatment of Laboratory Animals (Ministry of Science and Technology of the People’s Republic of China, Policy No. 2006 398). All biological experiments for JEV in this study have been approved by the Institutional Biosafety Committee (IBC) of the Yangzhou University, and performed in the biosafety level 2 facility of Yangzhou University, strictly following all biosafety management and regulation.

### Mouse studies

Three-week-old C57BL/6 mice were purchased from the Laboratory Animal Center of Yangzhou University and randomly divided into four groups (10 mice per group). Mice were respectively injected intraperitoneally with rGI(P0) [rescued with ppp(5′)A-RNA], rGI(P0) [rescued with m^7^G(5′) ppp(5′)A-RNA] or rGI-ΔDB2(P0) [rescued with m^7^G(5′) ppp(5′)A-RNA] at a dose of 10^3^ or 10^5^ TCID_50_ per animal, or mock-infected with equal volume of DMEM. All animals were monitored daily for death and clinical signs of disease.

### RNA *in vitro* transcription and virus recovery in cells

The JEV infectious clones TAR-rGI and TAR-rGIII were constructed in our lab(Li et al., 2022b). The JEV cDNA clones for RNA *in vitro* transcription were generated by cleavage linearization with restriction endonuclease *Bsu36I* (NEB). The full-length viral RNA was transcribed *in vitro* from the linearized JEV cDNA templates using the HiScribe T7 Quick High Yield RNA Synthesis Kit (NEB), and different 5′ cap structures of the viral genome were obtained by the addition of m^7^G(5′)ppp(5′)A cap analog or G(5′)ppp(5′)A cap analog (NEB). 2μg of transcript RNA was subsequently transfected into host cells seeded in a six-well plate with Lipofectamine MessengerMAX transfection reagent (Thermo Fisher Scientific) according to the manufacturer’s instructions. Culture supernatants of the transfectants exhibiting the typical cytopathic effects (CPE) of JEV infection were harvested and stored at -80℃ for further infection.

### Monocistronic and bicistronic reporters

The monocistronic reporter constructs used in this study were generated using the pGL3 vector backbone. The GI JEV 5’ UTR or GI JEV 5’UTR plus a 54-bp-long sequence in the 5’coding region of the C protein was respectively fused with the FLuc gene followed by either the GI JEV 3’UTR sequence or a short polyadenylated [poly(A)] tail (n=7) to construct monocistronic reporters, including JEV-5’UTR-FLuc, JEV-5’UTR-FLuc-3’UTR, JEV-5’UTR-cHP-cCS-FLuc and JEV-5’UTR-cHP-cCS-FLuc-3’UTR. Additionally, the monocistronic reporter HCV-5’UTR-FLuc with the FLuc gene flanked by an HCV IRES and a poly(A) tail was used as a positive control, and the HCV-IRES-Δdomain III-FLuc with the deletion of stem-loop III was used as a negative control. As above, the reporter RNAs were transcribed using T7 polymerase to contain either an m^7^G(5′)ppp(5′)A, G(5′)ppp(5′)A or ppp(5′) 5’terminal modification.

Bicistronic reporter constructs were generated using the pRL-SV40 vector as a backbone, including pRL-JEV-5’UTR-FLuc, pRL-JEV-5’UTR-cHP-cCS-FLuc, pRL-HCV-IRES-FLuc, and pRL-HCV-IRES-ΔdomainIII-FLuc. The upstream RLuc is flanked by an SV40 promoter and two stop codons, while the downstream FLuc is flanked by GI JEV 5’ UTR, JEV 5’UTR plus a 54-bp N-terminal coding region of the C protein, an HCV IRES or an HCV IRES-Δdomain III and a stop codon.

### Luciferase reporter assay

For the bicistronic reporter assay, the bicistronic pRL-JEV-5′UTR, pRL-JEV-5′UTR-cHP-cCS-FLuc, pRL-HCV-IRES or pRL-HCV-IRES-Δdomain III reporter plasmids were respectively transfected into HEK-293T cells for 24h. The cells were collected and lysed in 1×cell lysis buffer, after which the activities of RLuc and FLuc in cell lysates were detected in a luminometer (SuperMax 3000FL) using a dual-luciferase reporter assay system (Vazyme). For the monocistronic expression assay, a total 1μg of RNA transcripts of JEV-5′UTR-FLuc, JEV-5′ UTR-FLuc-3′UTR, JEV-5′UTR-cHP-cCS-FLuc or JEV-5′UTR-cHP-cCS-FLuc-3′UTR was respectively transfected into BHK-21 cells using Lipofectamine MessengerMax reagent (Thermo Fisher Scientific). At 12 hpt, the luciferase activity of FLuc was measured by the addition of FLuc substrate (Vazyme).

The *gaussia* luciferase assay was performed as previously described(Li et al., 2022a). Briefly, culture supernatants of cells transfected with JEV replicon RNA were harvested and cleared by centrifugation at 4°C. The luciferase activity was determined in a luminometer (LumiStation-1800) by mixing 20μL culture supernatant with 50μL reaction substrate Coelenterazine h (20μM, pH 7.2; Maokang Biotechnology).

### Immunoblotting and indirect immunofluorescence assay

Immunoblotting analysis was performed as previously described(Han et al., 2020). Briefly, whole cell extracts were lysed in ice-cold RIPA lysis buffer (Biosharp) with protease inhibitor (Roche). The supernatants of cell lysates were collected by centrifugation, separated by 10% SDS-PAGE, and further transferred onto a nitrocellulose (NC) membrane (Cytiva). The membrane was blocked with 5% skim milk in PBS and then respectively incubated with primary and secondary antibodies. The visualization of protein bands was performed using Tanon 5200 Multi chemiluminescent imaging system (Tanon). For indirect immunofluorescence assay, cells were fixed in 4% paraformaldehyde for 1 h, permeabilized with 0.05% NP-40 for 15 min and blocked with 5% bovine serum albumin (BSA) for 1 h at 37 ℃. Subsequently, the cells were incubated with primary antibody for 1h, followed by secondary antibodies for 1h. The nuclei were stained with 4,6-diamidino-2-phenylindole (DAPI) (Solarbio) for 15 min and the fluorescence images of cells were taken with a fluorescence microscope (Carl Zeiss).

### CRISPR-Cas9-based Genome Editing

DDX3-KO cells were generated using the CRISPR/Cas9 editing method. A gRNA targeting DDX3 was designed using E-CRISP (http://www.e-crisp.org/E-CRISP/designcrispr.html). The gene knockout cells with green fluorescence were sorted into 96-well plates by FACS and confirmed by immunoblotting and sequencing the genomic DNA.

### RNA pulldown assay

RNA pulldown assay was performed as described in the previous study(Han et al., 2020). In brief, BHK-21 or 293T cells were lysed with cell lysis buffer containing 200 U/ml RNasin (Beyotime), and centrifugation at 16,000 g for 10 min at 4℃ was performed to remove cell debris. The biotinylated RNA was synthesized using the T7 polymerase with the addition of 1.25 μL of 20 mM Biotin-16-UTP (Roche). The biotinylated RNA was heated to 90°C for 2 min in RNA folding buffer (10 mM Tris [pH7], 0.1M KCl, and 10 mM MgCl_2_) and the mixture was shifted to room temperature for 20 min to allow proper secondary structure formation. For the biotinylated RNA-binding assay, a reaction mixture containing 500 μg of cell extract and 2μg of biotinylated RNA was prepared. The mixture, at a final volume of 100 μL, was incubated in the reaction solution (5 mM HEPES [pH7.1], 40 mM KCl, 2 mM MgCl_2,_ and 1U RNasin) for 2 hours at 30°C and then incubated with 100 μL of Streptavidin Magnetic Beads (NEB) for 30 min at room temperature. The RNA-protein complexes were washed five times with reaction solution buffer. The RNA binding proteins were eluted by boiling the beads in 0.1% SDS and were analyzed by immunoblotting or mass spectrometry.

### GST Pull-down Assay

GST-DDX3, GST-PABP1, and GST proteins expressed in *E. coli* were purified with GSH Magnetic Agarose Beads (Beyotime). 10 μg purified GST-DDX3, GST-PABP1, or GST protein was incubated with HEK-293T cell lysates with the expression of PABP1-Myc or DDX3-Flag for 8 h at 4℃. The protein complexes were precipitated with GSH Magnetic Agarose Beads and detected with anti-GST, anti-Myc, and anti-Flag antibodies.

### RNA immunoprecipitation (RIP) followed by RT-qPCR (RIP-qPCR)

A total of 2×10^7^ BHK-21 cells were infected with rGI or rGI-ΔDB2 at an MOI of 0.01 or mock infected. At 12, 24 and 36 hpi, the culture medium was replaced with cold 1×PBS, and cell monolayers were irradiated with short-wave UV light at 5,700×100 μJ per cm^2^. After cross-linking, the cells were harvested by scraping and centrifugation prior to being lysed in RIPA buffer (50 mM Tris [pH 7.6], 150 mM NaCl, 1.0% NP-40, 0.5% sodium deoxycholate, 0.1% SDS). Cell lysate was then incubated 4 hours with either mouse anti-DDX3 antibody or mouse IgG-linked beads at 4℃. Subsequently, RNA captured by RIP was recovered using the Total RNA Extraction Reagent (Vazyme). The enrichment of JEV mRNA in each sample was assessed via RT-qPCR by the specific primers (Table supplement 4). The outcomes were calculated by normalization of RNA abundance in RIP samples (IgG and DDX3 mutant) to respective inputs.

### Polysome profile analysis

Cells were incubated with 0.1mg/ml CHX (Sigma) for 5 min at 37°C, and then lysed in polysomal extraction buffer (20 mM Tris-HCl [pH 7.5], 5 mM MgCl2, 100 mM KCl, 1% Triton X-100, 0.1 mg/ml CHX, 1×protease inhibitor cocktail, and 50 U/ml RNase inhibitor) for polysome fractionation by sucrose density gradient ultracentrifugation according to previous reports(Han et al., 2020). After centrifugation, each 500 μL fraction was collected and analyzed by UV absorbance at 254 nm. For analysis of proteins in polysomes, total proteins from each sucrose gradient fraction were precipitated with trichloroacetic acid (TCA) and analyzed by immunoblotting. To examine the distribution of ribosomal RNA in the gradients, total RNA was extracted with TRIzol (Vazyme) for quantitative RT-PCR analysis using specific primers targeting JEV and β-tubulin cDNAs (Table supplement 4).

### Secondary structure modeling

The secondary structure of JEV UTR was predicted with the Mfold software v 3.6 (https://www.unafold.org/mfold/applications/rna-folding-form-php) under the parameters of folding temperature at 37℃, an ionic condition of 1M NaCl with no divalent ions, and the VARNA program (https://varna.lri.fr/) was employed to modify the RNA structures.

### Statistical analysis

Statistical analysis was conducted using the unpaired Student’s *t*-test, or one-way analysis of variance (ANOVA) with Bonferroni’s post-tests for multiple-group comparisons. Animal survival analysis employed the Kaplan-Meier method to generate graphs and log-rank analysis was used to analyze the survival curves. *P* values less than 0.05 were considered statistically significant.

## Acknowledgements

This work was supported by the National Natural Science Foundation, China (grant no.32202886), the China Postdoctoral Science Fund (Grant No. 2022M722694), Natural Science Foundation of Jiangsu Province, China (Grant No. BK20220576), Jiangsu Co-innovation Center for Prevention and Control of Important Animal Infectious Diseases and Zoonoses, the Innovation and Entrepreneurship Team of Jiangsu Province (grant No. JSSCTD202224), and the Priority Academic Program Development of Jiangsu Higher Education Institutions (PAPD). Chenxi Li and Yanhua Li were supported by the Scientific Research Foundation of Yangzhou University and the “LvYangJinfeng Program” of Yangzhou City.

## Data availability

All data generated or analyzed during this study are included in the manuscript and supporting file.

---

**Figure 1-figure supplement 1. JEV infection was not restricted by the suppression of cap-dependent translation initiation.** (A-D) The puromycin incorporation assay of BHK-21, ST, DF-1 and C6/36 cells infected with 0.05 MOI JEV for 12, 24 and 36 h. (E and G) The puromycin incorporation assay at 12, 24 and 36 hpi of JEV in ST cells treated with 100 pmol eIF4E-specific siRNA (E) or 20 μM 4E2RCat (G). (F and H) The viral titers in the supernatants of BHK-21 cells treated with eIF4E-specific siRNAs (F) or 4E2RCat (H). ns, no statistical differences; Data are presented as mean ± standard deviation (SD) and statistically significant tested by Student’s *t*-test (F and H).

**Figure 2-figure supplement 1. Schematic diagram of reporters constructed for the identification of key elements for JEV cap-independent initiation.** (A) Schematic illustration of bicistronic constructs pRL-JEV-5′UTR, pRL-JEV-5′UTR-cHP-cCS, pRL-HCV-5′UTR and pRL-HCV-5′UTR-Δdomain III. (B) Diagrams of JEV monocistronic reporter constructs controlled by T7 promoter: HCV-5′UTR-FLuc, HCV-5′UTR-Δdomain III-FLuc, JEV-5′UTR-FLuc, JEV-5′UTR-cHP-cCS-FLuc, JEV-5′UTR-FLuc-3′UTR, JEV-5′UTR-cHP-cCS-FLuc-3′UTR and JEV-5′UTR-ΔSLA-FLuc-3′UTR. (C) Diagrams of JEV monocistronic reporter constructs with the deletion of each *cis*-acting element within 3′UTR.

**Figure 2-figure supplement 2. The immunofluorescence assay and replication characteristics of rescued viruses in BHK-21 cells.** (A) Immunofluorescence analysis of BHK-21 cells transfected with the JEV genomic RNA with 5′termini m^7^G(5′)ppp(5′)A or ppp(5′)A of WT or deletion mutants. (B) Plaque morphologies of the rescued viruses in BHK-21 cells, including rGI, rGI-ΔSLI, rGI-ΔSLII, rGI-ΔSLIII and rGI-ΔSLIV. (C) Growth kinetics of the WT and ΔSLI, ΔSLII, ΔSLIII, ΔSLIV mutant viruses in BHK-21 cells. Data are presented as mean ± SD from three independent experiments (C).

**Figure 2-figure supplement 3. Sequencing data of deletion mutant viruses. Related to Figure 2**.

**Figure 2-figure supplement 4. GLuc-based JEV replicon system for analyzing the translation and replication of viral genomic RNA.** (A) Schematic diagram of GLuc-based JEV replicon system. (B-D) GLuc activities in BHK-21 cells transfected with JEV replicon RNAs or mutants with 5’terminal modified by either m^7^G(5′)ppp(5′)A or ppp(5′). Data are the means ± SD of three independent experiments (C and D). The significant differences between m^7^Gppp(5′)A-rGI-GLuc and m^7^G(5′)ppp(5′)A-rGI-GLuc-ΔsHP-SL are labeled (^&&&^, p<0.001); The significant difference between m^7^Gppp(5′)A-rGI-GLuc and m^7^G(5′)ppp(5′)A-rGI-GLuc-NS5mut is marked (**^###^,** p<0.001); The significant difference between m^7^Gppp(5′)A-rGI-GLuc and m^7^G(5′)ppp(5′)A-rGI-GLuc-ΔDB2, or between ppp(5′)A-rGI-GLuc and ppp(5′)A-rGI-GLuc-NS5mut is marked (***, p<0.001).

**Figure 3-figure supplement 1. GLuc-based JEV replicon system for analyzing the translation of viral genomic RNA under the suppression of cap-dependent translation.** The BHK-21 cells treated with eIF4E specific siRNA or inhibitor 4E2RCat were respectively transfected with replicon RNAs: [m^7^G(5′)ppp(5′)A]-rGI-GLuc, [ppp(5′)A]-rGI-GLuc, [m^7^G(5′)ppp(5′)A]-rGI-GLuc-ΔsHP-SL to analyze the expression dynamics of GLuc. At different time points after transfection, the supernatants of BHK-21 cells were harvested and mixed with reaction substrate Coelenterazine h (20 µM, pH 7.2) to measure luciferase activity. Data are presented as the mean±SD of three independent experiments. The significant differences of m^7^Gppp(5′)A-rGI-Gluc-ΔsHP-SL between the treated (eIF4E-siRNA or 4E2RCat) and untreated (siNegative or DMSO) groups were labeled (***, p<0.001; ** p<0.01).

**Figure 5-figure supplement 1. Mass spectrometry identification of DDX3 and PABP1 as potential binding partners of JEV 3′UTR.** Red indicates matched B ions, blue indicates matched Y ions,grey indicates precursor ions.

**Figure 5-figure supplement 2. Generation of DDX3 knock-out BHK-21 cells.** (A) Schematic illustrating Cas9 inactivation of the hamster DDX3 locus. The 20-bp guide RNA target sequence is shown in blue, and the protospacer-adjacent motif (PAM) is shown in red.(B) Identification of the deletion within the DDX3 gene of DDX3-KO BHK-21 cells by DNA sequencing. (C) Analysis of DDX3 expression in WT and DDX3-KO BHK-21 cells by immunoblotting. (D) Cell proliferation of WT and DDX3-KO BHK-21 cells. ns, no statistical differences. Data are presented as mean ± SD from three independent experiments and tested by Student’s *t*-test (D).

**Figure 7-figure supplement 1. Identification of the secondary structures within UTRs bound by DDX3 and PABP1.** (A) Predicted secondary structures of DB2 and sHP-SL in 3′UTR and structures resulting from introduced mutations. Introduced substitutions are shown in red. (B) Predicted secondary structures of SLA in 5′UTR and structures resulting from introduced mutations. Introduced substitutions are shown in red.

